# Automated adjustment of metabolic niches enables the control of natural and engineered microbial co-cultures

**DOI:** 10.1101/2024.05.14.594082

**Authors:** Juan Andres Martinez, Romain Bouchat, Tiphaine Gallet de Saint Aurin, Luz María Martínez, Luis Caspeta, Samuel Telek, Andrew Zicler, Guillermo Gosset, Frank Delvigne

## Abstract

A lot of attention has been given to the understanding of microbial interactions leading to stable co-cultures, but the resulting technologies have been rarely challenged in dynamic cultivation conditions. In this work, substrate pulsing was performed to promote better control of the metabolic niches corresponding to each species, leading to the continuous co-cultivation of diverse microbial organisms. For this purpose, we used a cell-machine interface relying on automated flow cytometry, allowing to adjust the temporal profile of two metabolic niches according to a rhythm ensuring the successive growth of two species i.e., in our case a yeast and a bacterium. The resulting approach, called Automated Adjustment of Metabolic Niches (AAMN), was successfully employed for stabilizing both cooperative and competitive co-cultures. Additionally, AAMN can be considered as an enabling technology for the deployment of co-cultures in bioprocesses, demonstrated here based on the continuous bioproduction of p-coumaric acid. Taken altogether, the data accumulated suggest that AAMN could be used for a wider range of biological systems, but also to gain fundamental insights about microbial interaction mechanisms.

## Introduction

Being able to ensure the coexistence of several microbial species in the same cultivation device is a technological advance holding a lot of promises, both in terms of fundamental research (e.g., understanding how species can coexist and what are the ecological drivers for this coexistence^1234^), and applications (e.g., through the exploitation of division of labor for bioproduction^567^). The coexistence of microbial species is hampered by the competitive exclusion principle, but it is possible to overcome this principle in a number of cases by providing them access to alternative metabolic niches^8^. A simple way to diversify the metabolic niches is by pulsing nutrients into the cultivation device^9,10^. Even by pulsing only one type of nutrient (carbon source), it is possible to generate a temporal succession of metabolic niches through overflow metabolism and diauxic shift regulation, leading to the successive growth of the different species involved in the co-culture^8^. However, the determination of the amplitude and the frequency to be considered is not straightforward and depends, among other factors, on the global environmental conditions and the microbial species under consideration. Additionally, depending on the degree of cooperativity between the microbial species, environmental perturbation can lead to unexpected effects, such as increased stability for the most cooperative co-cultures or decreased stability for highly competitive co-cultures^1^.

In a previous work, we investigated the impact of environmental perturbations on the stability of a highly competitive co-culture involving *Saccharomyces cerevisiae* and *Escherichia coli* in a continuous cultivation device^11^. In a conventional metabolic niche where glucose was the main carbon source, the competitive exclusion principle led to progressive exclusion of the slow grower (*S. cerevisiae*) and the dominance of the fast grower (*E. coli*). However, we observed that, upon the diauxic shift on alternative substrates released upon overflow metabolism i.e., ethanol and acetate, *S. cerevisiae* exhibited transiently a higher fitness. Cross-feeding based on overflow metabolites is indeed known to promote community stabilization^1,12^. We then decided to exploit this metabolic niche by applying glucose pulsing at specific frequencies and amplitudes. For this purpose, we developed a model called MONCKS (for MONod-type Co-culture Kinetic Simulation) involving individual Monod-based models for each species. A cybernetic modeling approach was used to estimate the substrate utilization for each species in function of the appearance of the metabolic niches, with growth rate as the metabolic optimization target. In this way, we were able to simulate several scenarios for the coexistence of yeast and bacteria, and experimentally verified them. While being effective, this approach didn’t allow us to reach a co-culture with an equal number of individuals of both species. Additionally, the number of possible scenarios that can be simulated by MONCKS is too high to be tested experimentally.

The present work relies on the concept of ecological or metabolic niches i.e., the set of abiotic factors leading to the growth performance of a given species^13,14^, the maintenance of a single metabolic niche promoting the dominance of one microbial species over the others. To ensure the co-existence of two microbial species, it is thus necessary to alternate between two distinct metabolic niches at a frequency which is compatible with the growth capabilities of each species. Here, we adapted a cell-machine interface for automatically adjusting the transition between selected metabolic niches and ensuring the coexistence of different types of yeast-bacteria co-cultures exhibiting different mechanisms of interaction, an, approach called Automated Adjustment of Metabolic Niches (AAMN). The success of this operation is not straightforward since the metabolic niches can exhibit overlaps and variation in time i.e., niche expansion and contraction^11516^. Additionally, it is not easy to precisely define a metabolic niche, more specifically when complex microbial interactions are involved^17,18^. We then further expanded the MONCKS toolbox to directly integrate publicly available metabolic network models as a predictive tool for the optimization of the cell-machine interface. Indeed, several studies have pointed out that a metabolic niche can be captured more efficiently based on a quantitative biology approach using concept derived from metabolic flux analysis^19,20,17,16,18,21,15^. Based on these models, we were able to observe that metabolic niches overlapped for all the co-cultures involved in this work, but stability could be reached based on AAMN due to either metabolite cross-feeding or to the precise timing for the adjustment of metabolic niche. Finally, based on these observations, we successfully implemented AAMN for the continuous bioproduction of p-coumaric acid, and important building block for chemical synthesis^22^.

## Results

### 1. Automated Adjustment of Metabolic Niche (AAMN) as a generic strategy for the stabilization of cooperative and competitive co-cultures

The succession of two metabolic niches at an appropriate frequency could be used as a generic strategy for stabilizing the composition of co-culture^23,24,10,17,8^. Accordingly, we designed a cell-machine interface that automatically triggered the addition of substrate pulses based on the composition of the co-culture (**Figure 1a**). This interface comprises an automated flow cytometer (FC)^25^ that allows sampling the culture at regular time interval i.e., 15 minutes. The forward (FSC-A) and side-scatter (SSC-A) channels are used to discriminate between yeast and bacterial cells based on their size/morphology. In our case, control is applied for ensuring equal abundance of each species i.e., 50% of yeast and 50% of bacteria, during continuous cultivation. Based on this simple rule, pulse of carbon source compatible with one of the species is triggered when its abundance increases above 50%. We applied previously a similar approach for controlling phenotypically different subpopulations in mono-cultures based on a cultivation device called the Segregostat and involving reactive flow cytometry for promoting a specific population state^26,27^. To estimate the potential of our approach, called Automated Adjustment of Metabolic Niches (AAMN), to be generalized for diverse biological systems, we started the experiments based on two types of yeast-bacteria co-cultures differing at the level of their microbial interactions mechanisms.

**Figure 1:**
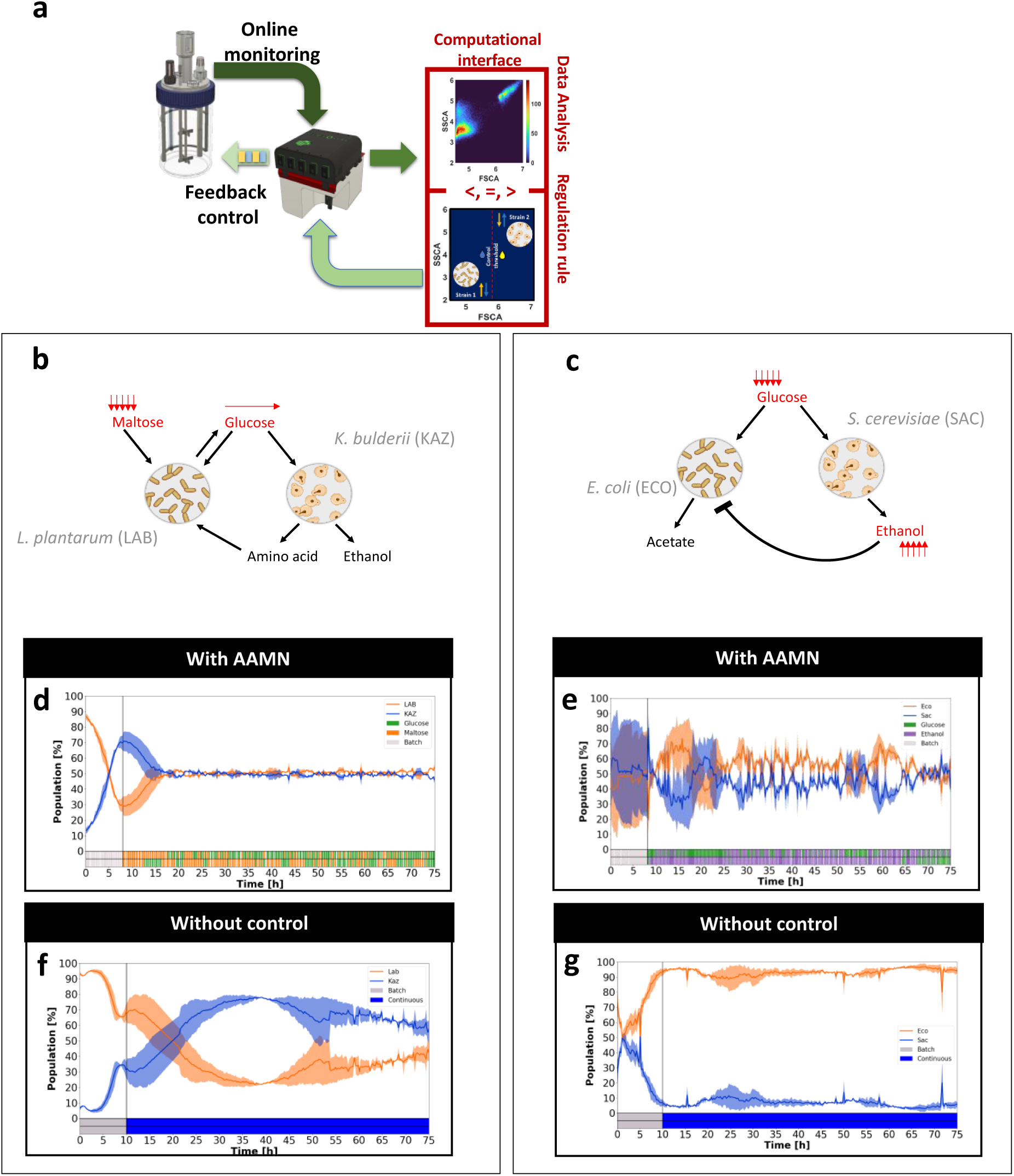
Automated reversion of metabolic niches can be used for stabilizing either cooperative or competitive co-cultures. **a** Cell-machine interface relying on automated FC allows for the control of co-culture composition (50%-50% for each species is selected here as a setpoint). Briefly, samples are taken automatically and are diluted before being analyzed by FC. Based on the analysis of the forward (FSC-A) and side (SSC-A) scatter signals, bacterial and yeast cells can be easily distinguished from each other. When the setpoint is not fulfilled, pulses of substrates (the nature of the substrate depends on the biological systems considered) are applied to restore equal abundance between the species. **b** Scheme of the metabolic interactions occurring between *Lactobacillus plantarum* (LAB) and *Kazachstania bulderii* (KAZ). **c** Scheme of the metabolic interactions occurring between *Escherichia coli* (ECA) and *Saccharomyces cerevisiae* (SAC). **d** Evolution of the number of cells (as determined based on automated FC) for a LAB-KAZ co-culture in a continuous cultivation device with AAMN (nutrient pulsing profile at the bottom of the figure is shown for n = 2 biological replicates). **e** Evolution of the number of cells (as determined based on automated FC) for a ECO-SAC co-culture in a continuous cultivation device with AAMN (nutrient pulsing profile at the bottom of the figure is shown for n = 2 biological replicates). **f** Evolution of the number of cells (as determined based on automated FC) for a LAB-KAZ co-culture in chemostat without AAMN. **g** Evolution of the number of cells (as determined based on automated FC) for a ECO-SAC co-culture in chemostat without AAMN. All the cultivations have been performed in duplicate (n = 2).

The first candidate is a co-culture involving *Kazakstania bulderii* (KAZ) and *Lactobacillus plantarum* (a lactic acid bacteria, LAB), a combination typically found in sourdough^28^ (**Figure 1b**). This type of co-culture is known to exhibit several commensalism interactions, the yeast providing resources (amino acids and nucleic acids) to the LAB^29,30^. During the implementation of AAMN for stabilizing KAZ-LAB, we also pulsed glucose to the co-culture. The Maltose present in the continuous feeding cannot be consumed by KAZ, but can be partially assimilated by LAB, one molecule of glucose being phosphorylated and assimilated by LAB, the other molecule being released into the extracellular medium^31,32^. The released glucose can then be utilized by KAZ, leading to a double cross-feeding system that is known to promote co-culture stability^33,34,35^. The second co-culture considered involves *Saccharomyces cerevisiae* CEN.PK 113-7D (SAC) and *Escherichia coli* W3110 (ECO) (**Figure 1c**). This co-culture exhibits more complex competitive/commensalism interactions, notably all the excretion of overflow and/or fermentation metabolites (i.e., mainly ethanol and acetate) that can act as alternative carbon sources and promote cross-feeding, competition and even exclusion by inhibitory effects. Taken altogether, these alternative metabolic pathways will determine the establishment of the proper metabolic niches. We previously demonstrated that it is possible to stabilize co-cultures (20% SAC; 80% ECO) based on periodic pulsing of glucose^11^. Under these conditions, *E. coli* exhibited the highest fitness when glucose is high, but *S. cerevisiae* exhibited more fitness at low glucose concentration and upon reassimilation of acetate and ethanol. The pulsing frequency was predicted based on a cybernetic model and several scenarios, with different level of co-cultures stability, were predicted and experimentally verified^11^. Applying environmental perturbations at given frequencies has been shown to be effective for the stabilization of co-cultures^9^, as well as in more complex microbial communities^10^. However, finding the appropriate perturbation or pulsing frequency is not straightforward. We then focused on this aspect by using our cell-machine interface to verify whether the appropriate pulsing frequency can be reached automatically for driving equal composition during the whole continuous cultivation. Taken altogether, our data point out that co-culturing LAB-KAZ (**Figure 1d**) and ECO-SAC (**Figure 1e**) based on the AAMN approach leads to the stabilization of the co-cultures, by comparison to continuous cultivation performed without control (**Figure 1f-g**). However, we can see that it is more difficult to stabilize ECO-SAC (**Figure 1e**) that LAB-KAZ (**Figure 1d**), suggesting that the alternance of metabolic niches is not straightforward in this case. In the next section, metabolic and dynamic modelling will be considered to simulate the succession of metabolic niches and its impact on co-culture dynamics.

### 2. Full reversion of metabolic niches is needed for the stabilization of competitive co-cultures

We have seen in the previous section that the LAB-KAZ and ECO-SAC co-cultures display different dynamics. For LAB-KAZ, we observe a cooperative behavior where co-culture stabilization can occur during long-term continuous cultivation, even without control (**Figure 1f**, co-culture tends to equal abundance until the end of the experiment) or is reached even more quickly upon applying the AAMN approach (**Figure 1d**). The picture is totally different for ECO-SAC, with a strong competitive behavior reported. This behavior can be clearly observed during continuous cultivation (**Figure 1g**) and can also be observed upon applying AAMN (**Figure 1e**), where microbial species tend to escape control as they have the metabolic capability to contend for all metabolic niches. To understand what drives either competitive or cooperative behavior, we considered Elementary Mode (EM) analysis of the central metabolism derived from existing genome-scale and other metabolic models for the different species involved (**Figure 2a-b, Supplementary Notes 1 and 2**) to quantitatively characterize the degree of competitivity θ between strains (**Figure 2c**). θ depends on the metabolic niches (MNs) available for the growth of each species. As an example, if a substrate can be used by both species in the co-culture, it will lead to high competitivity and, accordingly, a θ value close to or equal to 1. On the other hand, if another substrate can be used by only one of the species, this will lead to the appearance of an exclusive MN promoting the growth of this species only, ultimately leading to a reduction of θ. We then applied this approach to the analysis of the metabolic niches that have to be expected during the continuous cultivation experiments^21,18,36^. In an ideal case, we could exploit totally exclusive niches (θ = 0) for each strain and alternate between them at a given rhythm adjusted to the desired mean growth rates. However, in reality, metabolic niches commonly overlap, increasing the competitivity and reducing the controllability of the system over time^37,19^.

**Figure 2:**
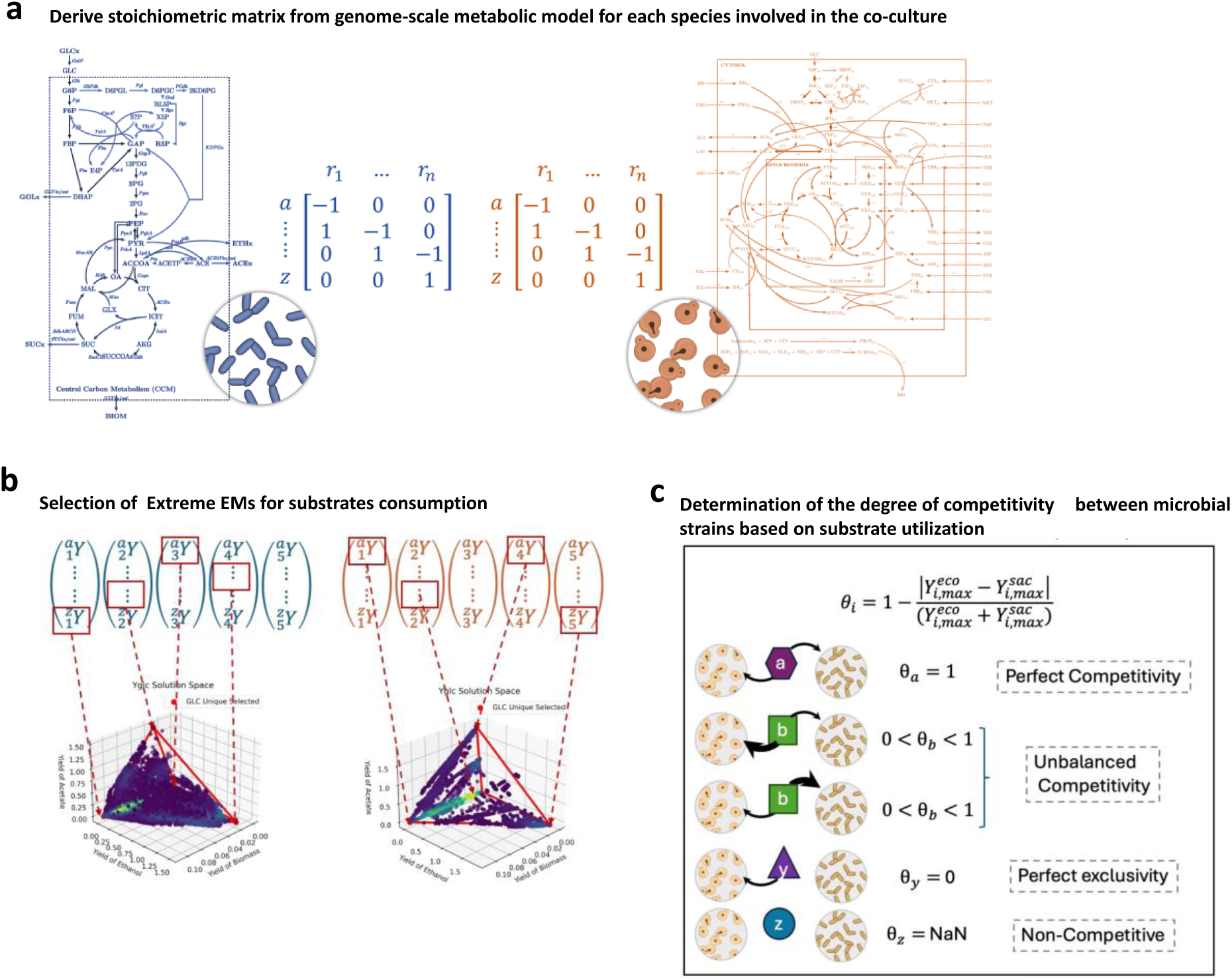
A framework for computing competitivity between strains involved in co-cultures. **a** Genome-scale models for ECO and SAC are processed to generate a stoichiometric matrix for each species. **b** Elementary Mode (EM) analysis is performed on each stoichiometric matrix to identify the major pathways involved in biomass generation. Yield coefficients i.e., Y_i_, are then computed for each species and substrates. **c** For each metabolites/substrates available in the extracellular medium, the degree of competitivity θ between strains can be computed based on the respective Y_i_. Different scenarios are shown.

In short, we determined the most relevant metabolic pathways for each species in co-culture pairs based on genome-scale metabolic network and EM analysis (**Figure 2a-b, Supplementary Note 2**). Based on this approach, we were able to determine the metabolic space for each microbe resulting from the metabolic flux distributions with the major substrate consumption capabilities used for biomass generation based on by-products utilization (**Figure 2b**). The results can be displayed in a yield convex solution space for three different potential substrates (Figure 2b), where each dot represents an EM and its position the biomass yield (Y_x/s_) for any specific substrate. The most extreme EMs positions for each substrate can be selected to construct a niche polygon (NP) which contains all the MNs that can be used by each strain. Comparing the vertices of the polygons of each strain is then easy to visualize The MN preference for each strain can be easily visualized based on the vertices of the polygons and can be used for computing the competitive advantage for each of the strains in the co-culture (**Figure 2c**). These polygons are multidimensional and contain the total capabilities of a strain. However, they can also be simplified to a selected active set of metabolic pathways, resulting from only accounting for the solutions concerning the existing environmental substrates where the microbial species are being cultured (see **Figure 3a-b** for ECO-SAC)^18,13^. For better visualization, in the case of LAB-KAZ, two different NPs were constructed i.e., one for the main carbon sources, and the other for the amino acids (**Supplementary Note 2**). For the carbon source, the analysis of MN points out a clear overlap between LAB and KAZ. However, in this case, competitive exclusion can be avoided based on maltose feeding and glucose pulsing during AAMN experiment and the resulting glucose cross-feeding. For the amino acid, the NP revealed that KAZ produce glycine and isoleucine that can enter in a cross-feeding metabolic flux with LAB relieving its auxotrophy. In the case of KAZ-LAB there is an increased degree of cooperation due to the auxotrophies from LAB and the KAZ substrate inaccessibility which increases the stability of the co-culture, even in the face of the GLC metabolic overlap that can be observed based on the NPs. In other words, the temporal generation of suitable metabolic niches for each species is ensured by the mutual cross-feeding provided by the amino acids naturally released by the yeast and the maltose pulsed during continuous cultivation. As a result, the AAMN leads to a very smooth control of the co-culture composition over time (**Figure 1f**).

**Figure 3:**
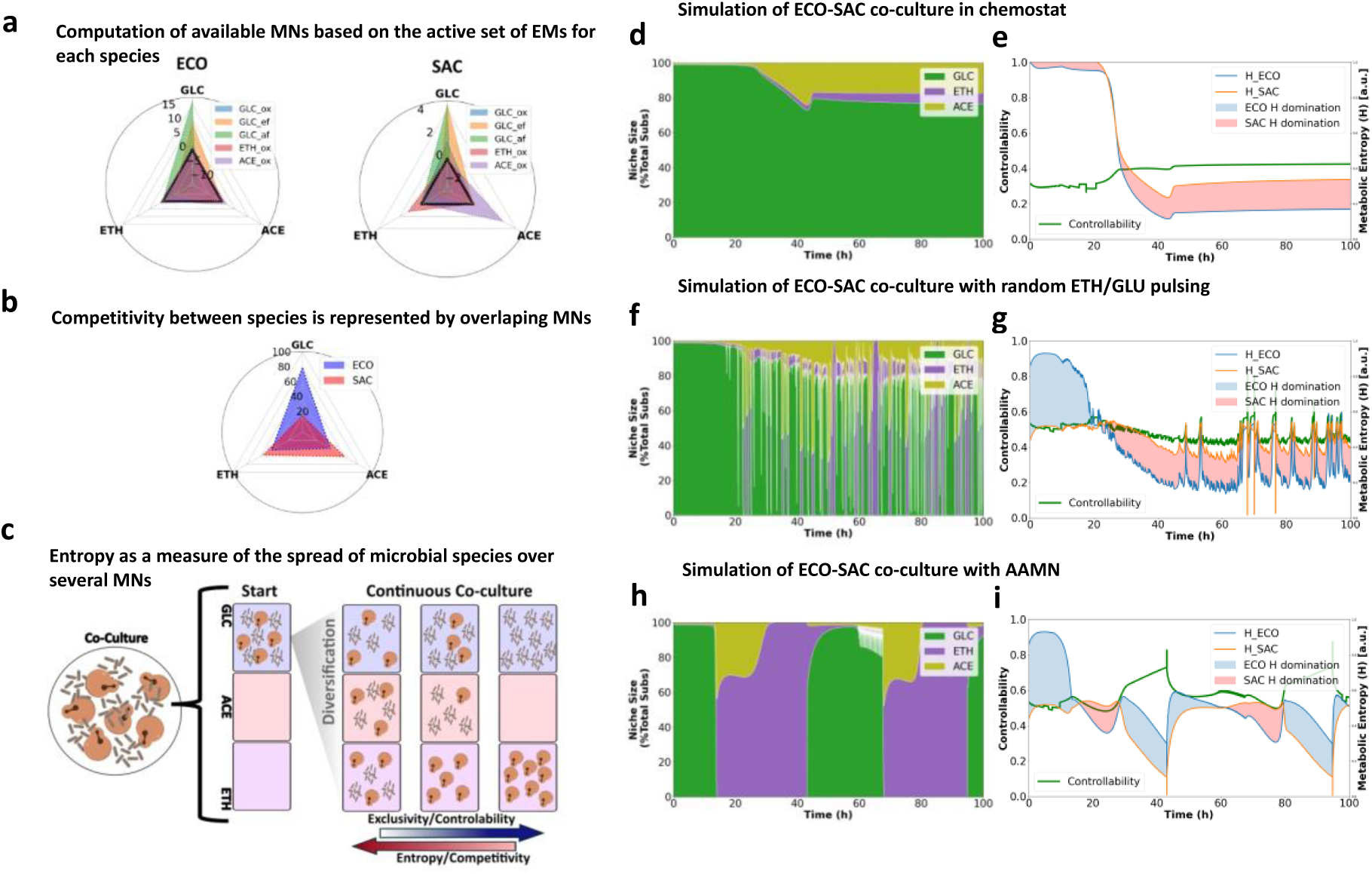
Full reversion of metabolic niches is needed for stabilizing competitive ECO-SAC co-cultures. **a** Metabolic niche polygon representing the utilization of various substrate by the co-cultured species i.e., here ECO and SAC. These niche polygons have been computed based on the biomass yields Y_x/s_ obtained from the metabolic pathways contributing the most to growth based on EMs analysis. **b** Overlap between two niche polygons can be used for estimating the degree of competitivity between two microbial species, since the corresponding MNs are not exclusive in case of overlap. **c** Information entropy can be used for estimating the dynamic dispersion of one species to the mobilization of several MNs. When the computation is performed for two species, the comparative analysis of the two entropies can be used for determining either controllability based on te exclusivity of the MNs, or to compute the competitivity between species. **d** Simulation of the evolution of the MNs size for the ECO-SAC co-culture growing in chemostat. **d** Evolution of the controllability and entropy computed from the comparative MN repartition between ECO and SAC in chemostat. **f** Simulation of the evolution of the MNs size for the ECO-SAC co-culture in continuous mode with random substrate (ETH and GLU) pulsing. **g** Evolution of the controllability and entropy computed from the comparative MN repartition between ECO and SAC in a continuous cultivation device with random ETH/GLU pulsing. **h** Simulation of the evolution of the MNs size for the ECO-SAC co-culture in continuous mode where ETH and GLU are pulsed according to AAMN. **i** Evolution of the controllability and entropy computed from the comparative MN repartition between ECO and SAC in a continuous cultivation device with ETH/GLU pulsing performed based on AAMN (Abbreviation used: GLU: glucose; ACE: acetate; ETH: ethanol).

For ECO-SAC, the NP revealed that the metabolic niches for each species are all overlapping (**Figure 3a-b**), explaining the competitive behavior observed during the experiments (**Figure 3c**). In this case, this overlap is not compensated by cross-feeding. Indeed, even if overflow metabolites (ACE and ETH) are released, they are not mutually exclusive for one of the species, further contributing to the system destabilization as bacteria and yeast growth rates are significantly different in all metabolites. For computing the stability of the co-cultures, we would of course use the index θ. However, this index allows to compute the competitivity of the strains in fair and stationary conditions only. We then decided to set-up a quantitative approach for determining the dynamics of cooperativity/competitivity between microbial species in co-culture. For this purpose, we derived parameters from the computation of information entropy. Information entropy is frequently used for computing the degree of phenotypic heterogeneity of mono-culture based on the spread of the phenotypes distribution^38,39,27^. We use a similar approach for capturing the spread of one microbial species over the utilization of several MNs (**Figure 3c**). Based on the computation of the respective entropies related to MN utilization for each species, it is then possible to derive parameters about the degree of cooperativity/competitivity between species (**Supplementary Notes 2, 3 and 4**). The entropy is based on the estimation of the probability for a substrate to be consumed by a specific strain based on the simulated instantaneous consumption rates, therefore providing a more dynamic view of the competition rather than the previously formulated index θ. Furthermore, measuring the relative entropy exhibited by each strain in the co-culture allows us to determine how well the substrates are being distributed along the strains, and therefore, the exclusivity of substrate utilization and the competitivity. Additionally, this dynamic approach allows us to calculate how efficient pulses or perturbations would change the systems composition i.e., its controllability. We then used these proxies in different co-culture scenarios (**Figure 3d-i**). It has been observed that AAMN is very important in the case of ECO-SAC co-cultures for continuously adjusting the metabolic niches during co-culture. The precise adjustment of the metabolic niche based on substrate pulsing seems thus a very important element for stabilizing competitive co-cultures. To challenge this hypothesis, we conducted dynamic simulation of ECO-SAC co-cultures based on different cultivation scenarios i.e., either chemostat (**Figure 3d-e**), random pulsing cultivation (**Figure 3f-g**), or AAMN (**Figure 3h-i**). For this purpose, we used a previously developed dynamic modeling toolbox, called MONCKS^11^, complemented with the main metabolic pathways identified based on EM analysis. As expected, the dynamic simulations pointed out stabilization in the case of the cultivation conducted under AAMN, whereas population collapse was observed in the case of chemostat or random pulsing cultivation (this last scenario was further experimentally validated, **Supplementary Note 3**). These results point out that, even if the NPs reveal strong niche overlap, AAMN can be used for reverting the MNs at a specific frequency, leading to the stabilization of the co-culture^13^ (**Figure 3g-h**).

### 3. Engineering cooperativity in continuous co-cultures of *E. coli* and *S. cerevisiae*

Whereas the AAMN approach can be used for stabilizing co-cultures exhibiting competitive behavior, the results pointed out a population oscillatory profile (**Figure 1e**) which is maybe not convenient to exploit for application like bioproduction, where a more stable co-culture profile is expected^40^. Accordingly, we decided to implement a separation of carbon sources in an attempt to promote a cooperative behavior between *E. coli* and *S. cerevisiae*^54,16^. For this purpose, we used an *E. coli* strain defective in active glucose transport^42,43^, and the alternative pulsing of glucose an xylose for generating metabolic niches promoting the growth of *S. cerevisiae* (SAC*) and *E. coli* (ECO*) respectively. We then adapted the metabolic fluxes for these strains and recomputed the competitivity index θ. As expected, θ decreased, and increased the controllability of the system from 9.1 % to 28.3 % (**Supplementary note 2**). The ECO* strain exhibits deletion of components of the phosphotransferase system (PTS)-dependent sugar transporters, as well as non PTS-transporters^42,43^. As a results, this strain exhibited reduced glucose uptake and acetate excretion, allowing the yeast to grow mainly on glucose during co-cultivation. Xylose was considered as a second carbon source for ensuring the co-existence of the ECO* strain. This separation of carbon source is expected to provide more control over co-culture composition^56,40^. We adapted the metabolic fluxes accordingly and performed the EM analysis for defining the NP for the two microbial species (**Figure 4a**). These computations were complemented by dynamic simulations of the ECO*-SAC* co-culture based on MONCKS (**Figure 4b-d**). We first conducted the simulations by adjusting the uptake rate of glucose and xylose by ECO*-SAC*. In a first, simulation, we considered that the rate of glucose uptake and xylose was equivalent. While this computation should lead to the stabilization of the ECO*-SAC co-culture, simulation pointed out that the co-culture was destabilized after approximately 40 hours of cultivation (**Figure 4b**). Interestingly, this destabilizing effect was independent of the ratio between glucose-xylose uptake rates (**Figure 4d**). Furthermore, we see that the calculated entropy and controllability were very similar. The latter was the result of the appearance of an unexpected and uncontrolled MN involving acetate (ACE). Indeed, acetate is produced by *E. coli* upon overflow metabolism and can be reassimilated by both species upon adaptation in continuous cultures, leading to new MNs contributing to the destabilization of the co-culture (**Supplementary notes 3 and 4**). However, the ECO* strain is defective in the PTS are is not able to produce ACE due to its reduced GLU consumption rates^42,43^. However, they are still capable of ACE production if the pyruvate and phosphoenolpyruvate node are saturated, and that could be the case arising from GLC and XLY co-utilization. We then studied the possibility minimizing the ACE MN by reducing the EMs ACE Yield in the models and reprocessed dynamic simulation based on MONCKS (**Figure 4f**) which, this time, predicted co-culture stabilization based on AAMN. This result is important because it point out that the direct separation of carbon sources is not necessarily an effective approach for stabilizing co-culture, if cross feeding and other unexpected niches are not considered for the control strategy. Since in some cases, like the one considered here, consumption of alternative carbon source can lead to the emergence of unwanted MNs, further destabilizing the co-culture. These results point out that our simulation toolbox could be generalized for probing different metabolic engineering strategies aiming at stabilizing co-culture. In the next section, we’ll challenge experimentally the utilization of the ECO*-SAC* co-culture for the continuous bioproduction of p-coumaric acid.

**Figure 4:**
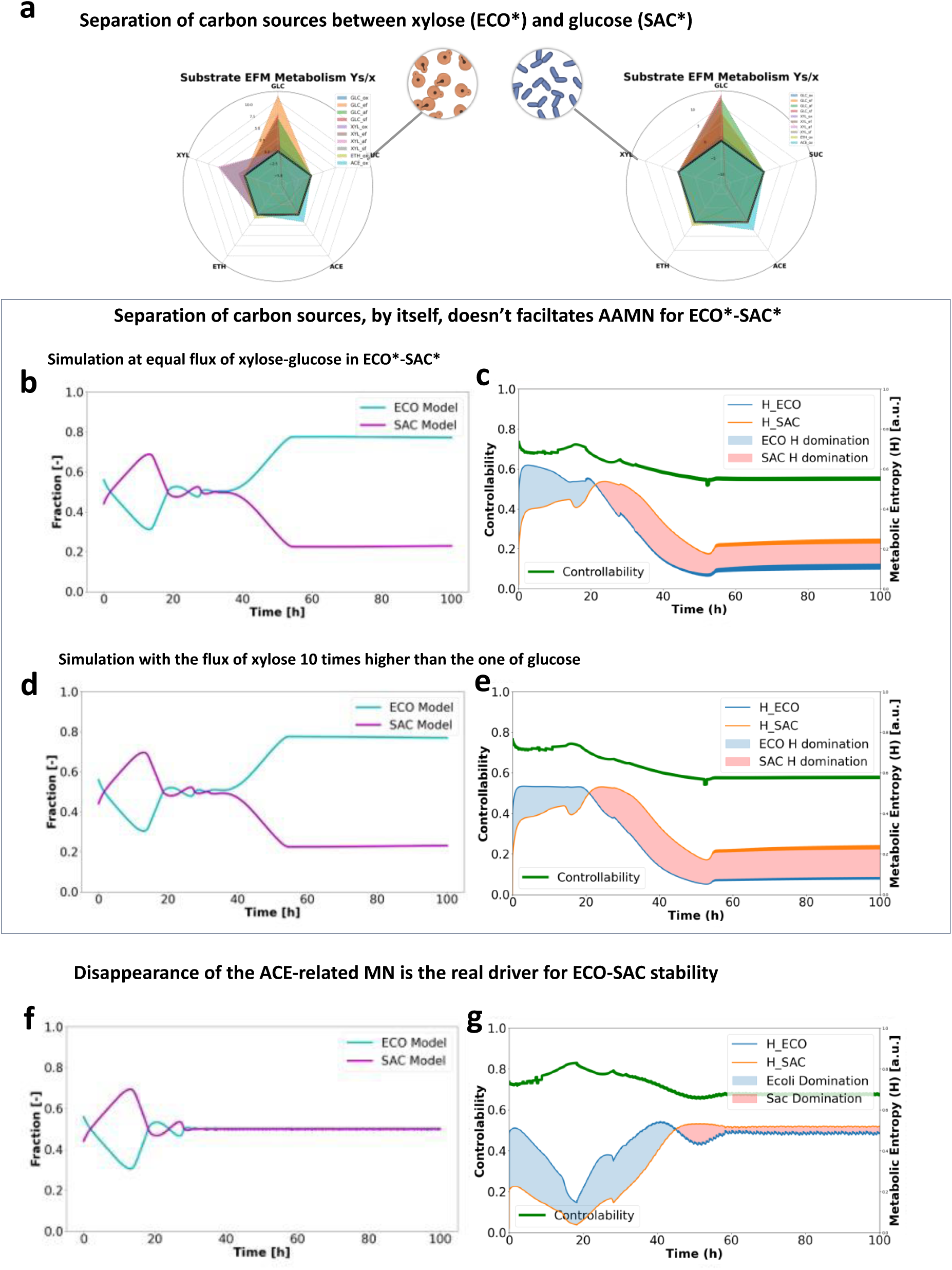
EM analysis and MONCKS predict that the suppression of the ACE-related MN promotes cooperativity in ECO-SAC co-cultures. **a** Metabolic niche for ECO* and SAC* computed from EM analysis of genome-scale metabolic models. The values represent the biomass yield Y_x/s_ for the different EMs, corresponding here to the different MNs. **b** Dynamic simulation based on MONCKS and showing the evolution of the fraction of ECO* and SAC* in AAMN continuous cultures at a dilution rate D = 0.1 h^-1^ with controlled additions of GLU and XYL (0.4 grams per pulse), GLU and XYL ECO maximum consumption rates with equal values **c** Evolution of the controllability and entropy of the in AAMN continuous co-culture operated with an equal feed GLU and XYL maximum consumption rates. **d** Dynamic simulation based on MONCKS and showing the evolution of the fraction of ECO* and SAC* in AAMN continuous cultures at a dilution rate D = 0.1 h^-1^ with controlled additions of GLU and XYL (0.4 grams per pulse), GLC ECO maximum consumption rates 10 times smaller than the ones for XYL. **e** Evolution of the controllability and entropy of the in AAMN continuous co-culture operated with XYL rates ten times bigger than the ones of GLU. **f** Dynamic simulation based on MONCKS and showing the evolution of the fraction of ECO* and SAC* in AAMN continuous cultures at a dilution rate D = 0.1 h^-1^ with controlled additions of GLU and XYL (0.4 grams per pulse), with reduced ACE and ETH production yields ten times. **g** Evolution of the controllability and entropy of the AAMN continuous co-culture operated in reduced ACE and ETH overflow metabolism. (Abbreviation used: GLU: glucose, XYL: xylose, ACE: acetate)

### 4. AAMN is effective for promoting division-of-labor and coumaric acid production by ECO*-SAC* co-culture

One of the key benefits of using co-cultures instead monocultures is their ability to enhance metabolic capabilities of the system, based on the division-of-labour principle^5,17,44,41^. Accordingly, we decided to adapt our potentially cooperative ECO*-SAC* co-culture (**Figure 4d**) for the continuous bioproduction of p-coumaric acid. ECO* and SAC* were then further engineered to produce tyrosine and its bioconversion to p-coumaric acid respectively (**Figure 5a**). We first carried out chemostat experiments where different xylose/glucose ratio were considered as co-feeding (**Figure 5b**). We can see that, upon reducing the amount of xylose in the feed, stabilization of the co-culture naturally occurs after 60 hours of cultivation, without the need for AAMN. This observation confirms that the separation of carbon sources can promote cooperative behavior in ECO*-SAC* co-cultures. We then considered AAMN for enhancing the stabilization of the co-cultures (**Figure 5c**). Upon AAMN, a partial stabilization of the co-cultures was observed readily after approximately 20 hours of cultivation. However, we can also see that the cooperative behavior for the ECO*-SAC* is not as elevated as the one reported for LAB-KAZ (**Figure 1d**) or the one predicted by MONCKS (**Figure 4d**). On the other hand, AAMN allowed for the stabilization of the ECO*-SAC* segregostat co-culture, resulting in a 3.5-fold improvement in p-coumaric acid yield when compared to chemostat cultures (**Figure 5d**).

**Figure 5:**
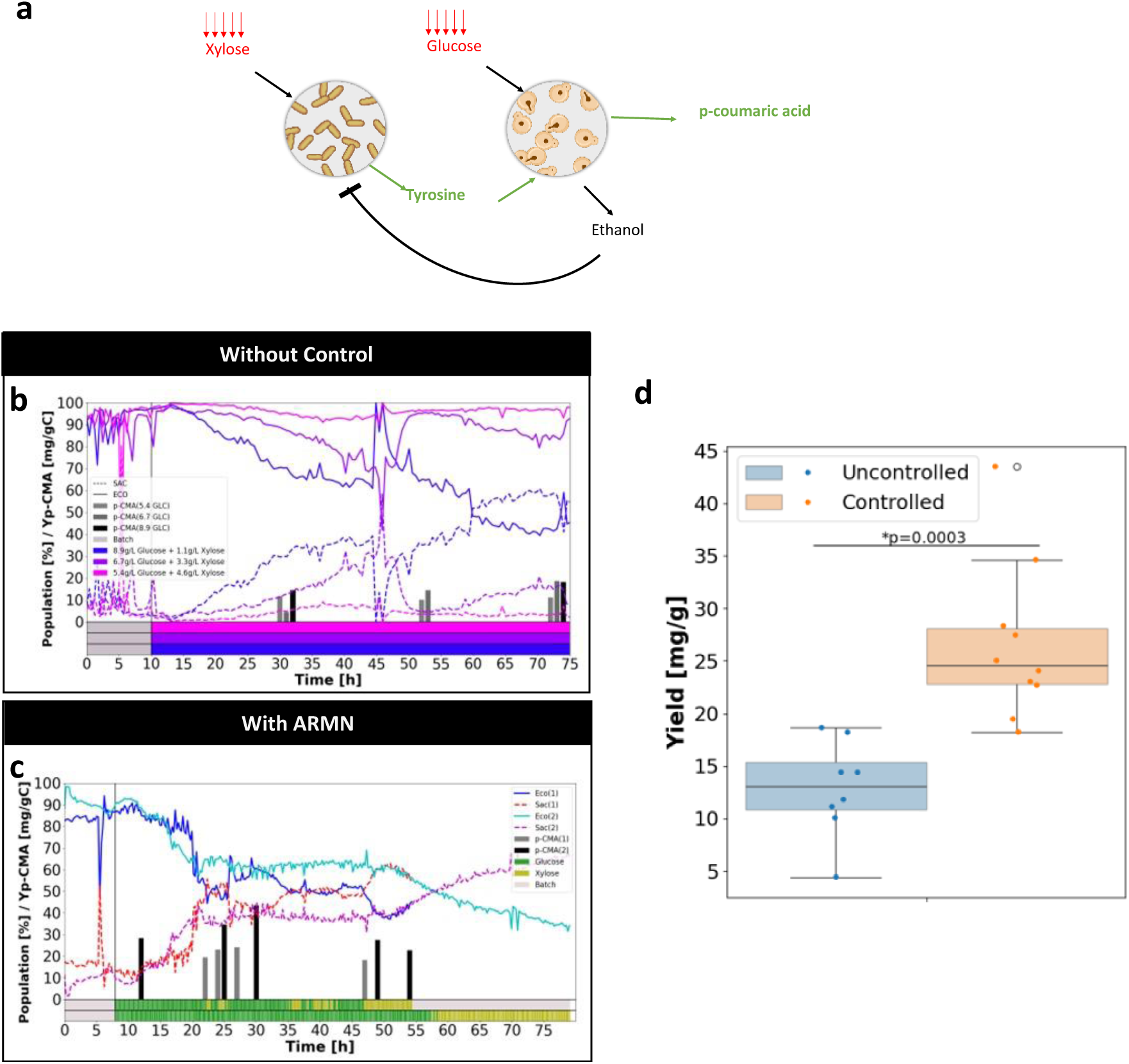
AAMN allows for the stabilization of ECO*-SAC* co-culture engineered to produce p-coumaric acid. **a** Scheme of the metabolic interactions taking place in the context of the ECO*-SAC* co-culture. **b** Evolution of the number of cells (as determined based on automated FC) for the ECO*-SAC* co-cultures in chemostat with variable xylose/glucose ratios in the feeding. **c** Evolution of the number of cells (as determined based on automated FC) for an ECO*-SAC* co-culture in a continuous cultivation device with AAMN in duplicate (nutrient pulsing profile at the bottom of the). **d** Comparison of p-coumaric acid yields between continuous cultures performed with and without AAMN, Yields were calculated from samples from all experiments after 24 hours, all samples from the AAMN were clustered to make the controlled type box plot (sample size = 10) and compared to all uncontrolled samples boxplot (sample size = 8), and ANNOVA single tail test was performed and revealed the significant difference between both populations with a p value of 0.0003 .

Several explanations can be advanced at this level. First, the toxicity of the product i.e., p-coumaric acid, can impact the stability of the co-cultures. Second, the theoretical destabilization we found with MONCKS in the previous section, by the apparition of unwanted or uncontrolled MN such as acetate or ethanol could impair AAMN. Third, cellular co- and/or auto-aggregation was observed based on automated FC profiles (**Supplementary note 5**). Aggregation has direct impact on co-culture stability since it has been previously reported that cell auto- or co-aggregation can lead to collective elimination in some ecosystems^45^. Also, auto-aggregation of yeast cells have been previously reported as decreasing the performances of bioethanol production upon bacterial contamination^46^. However, the FC data didn’t point out any difference in the intensity of cell aggregation between the different co-cultures i.e., either cooperative and competitive, suggesting that product toxicity and/or uncontrolled MNs could be the main driver for co-culture destabilization in the case of ECO*-SAC*.

## Discussion

Cell-machine interfaces, relying either on microfluidics^47,48^ or FC^27,49^ can be used for manipulating gene expression in individual cells within population, leading to the acquisition of data for characterizing gene circuits^50,47^ or, in a more applied perspective, to improved bioproduction^51,52,26^. Similar approach has been recently applied for manipulating co-culture composition over time^53,54^. In this work, we propose a generalizable approach, relying only on the use of nutrient pulsing, for adjusting the metabolic niches and strain co-existence. Based on our AAMN approach, we were able to control 3 different types of yeast-bacteria co-culture, including the ones exhibiting strong competitive behavior. Synthetic biology provides a lot of different technologies aiming at controlling co-cultures and/or microbial communities^23,55,56^. However, most of the technologies proposed so far come at a high metabolic load for the hosts^57,58^ or with harsh consequences for the targeted species e.g., cell lysis^59^. As an example, systems involving either toxin-antitoxin or bacteriocins seems very effective^60^, but results in cell lysis for controlling the dominance of one of the species. For applications, like bioprocessing, cell death and lysis must be avoided, since it can lead to loss of resources and productivity, as well as issues related to downstream processing operations. Unwanted cell lysis can be prevented based on the use of other technologies, such as the utilization of Quorum Sensing circuits, for the control of the cell density of the different species involved^61,62,63,64^. However, the signaling molecules used for probing and controlling cell density (e.g., AHL) can be diluted upon scaling-up, further impairing the transposition of the system to industrial applications. It is also possible to dynamically control gene circuits based on light (optogenetics)^51,49^, and this technology has already been used to controlling gene circuits for the stabilization of co-cultures^53^. Again, the utilization of light as an actuator for cell population control is difficult to up-scale, due notably to the limit of transmission of light and dense and large volume of cell suspension^65^. Here, we show that non-destructive strategies, like the separation of carbon sources, and the AAMN can be used for promoting cooperative behavior in ECO-SAC co-cultures. We also show that the timing in the adjustment of the MNs is critical for stabilizing co-culture composition. Accordingly, such approach requires the precise quantification of the related MNs. It is now possible to define more precisely the MNs based on the utilization of genome-scale metabolic models^19,17,21,18,15^. We carried out EM analysis based on the metabolic networks of the 6 natural of engineered species considered in this study. Surprisingly, we find out that only a limited set of EMs (i.e., between 3 and 5) are needed for recapitulating co-cultures dynamics (except in the specific case of amino acid exchange for LAB-KAZ where more than 20 EMs were involved) based on the use of the MONCKS toolbox^11^. Taken altogether, the experimental and numerical tools developed in this work can be used as a general approach for capturing microbial co-culture dynamics and enabling the development applications, such as continuous bioprocesses.

## Methods

### Strains and plasmids

The different co-cultures were done with different bacterial/yeast composition. The first system considered was *Escherichia coli* W3110 (ECO) and *Saccharomyces cerevisiae* CENPK *pchi-eGFP* (SAC). The second system was composed of *Lactobacillus plantarum* (LAB) and *Kazachstania bulderi* (KAZ) and the last couple tested was composed of *Escherichia coli* WHIC^66,42^ transformed with two plasmids i.e., *pJLBaroGfbr* and *pTrcTyrCpheACM* (ECO*)^67^ and *Saccharomyces cerevisiae* PTA 408 *+pCA17* (SAC*) for the production of p-coumaric acid (p-CA) from glucose (GLU)^68^.

### Media composition

Verduyn media was used for all cultivations involving *E. coli* and *S. cerevisiae*. MRS medium complemented with a vitamin solution (adapted MRS) was used for the cultivation of *L. plantarum* and *K. bulderi.* The compositions of these media can be found in **Supplementary note 6**. Both media were complemented with carbon sources for pre-cultivations and cultivations according to the detailed experiments. For pre-cultivations glucose 20 g/L was used for ECO and SAC, maltose-glucose 7.5 g/L each for LAB and KAZ, and finally glucose-xylose 5 g/L of each for ECO* and SAC*. Ampicillin (50 µg/L) was sterilized by filtration (0.2 µm) and added for ECO* and SAC* pre-cultivations for plasmid maintenance. Tetracycline (10 µg/L) was also sterilized by filtration (0.2 µm) and added for ECO*. The pre-cultivations were all performed at 30°C for yeasts and 37°C for bacteria and at 150 RPM overnight (16 hours) in non-baffled flasks.

### Chemostat and Segregostat cultivations

The cultivations were done all at least in duplicate in lab-scale stirred bioreactors (Bionet twin) with a working volume of 1 L and at an initial OD600 of 0.1 and with the corresponding medium (Adapted MRS or Verduyn). A batch phase was before continuous cultures or controlled cultures was used for biomass propagation. These batch phases lasted between 8 to 10 hours with the microorganisms growing in the corresponding medium complemented with carbon sources mix (same as for the precultivations).

Growth data and event fractions were collected based on online FC during the experiments with AAMN^69,70^. Briefly, it consists in an automatic system that takes sample every 15 minutes from the bioreactor, analyses the samples in a flow cytometer (BD Accuri C6plus, Biosciences) and an actuator pulsed different substrates by the means of peristaltic pumps activated on basis of the FC data and a custom MATLAB script. In this case, the actuator was pulsed based on the observed distribution between yeasts and bacteria (**Figure 1**). The percentage of these two populations activated the addition of one or two carbon sources accordingly. The carbon sources were provided by the feeding at same concentration than during batch phase for Chemostat cultivations.

*E. coli* and *S. cerevisiae* were cultivated at 1000 RPM and 1 VVM with Verduyn medium at a pH of 6.8. The inoculation population ratio for Chemostat cultures was 1:1 ECO/SAC (OD600/OD600.), and for the AAMN experiments it was 1:10 ECO/SAC. The latter was done to allow at the end of batch section have proportions near 50% in terms of events fraction and therefore allow control to take actions near the threshold set for the control rule in this case 50 %. In chemostat, the feed was set to be 10 g/L of glucose. For AAMN experiments, the feeding was done with Verduyn medium with no supplemented carbon source, and pulses were set to be either GLU or ETH each 15 minutes accordingly to the exceeding populations of *SAC* or *ECO*, respectively. Dilution rate was set to be 0.1 h^-1^ for both chemostat and AAMN cultivations, while pulses were set to add 0.4 grams of carbon source per pulse of either glucose or ethanol.

*L. plantarum* and *K. bulderi* were cultivated at a stirring frequency of 400 min^-1^, an air flow rate of 0.2 VVM with adapted MRS medium at pH of 5.6. The population ratio for inoculation was 1.85:1 LAB/KAZ. The feeding rate was 0.1 h^-1^ after the batch phase to reach around 50% of each strain. In chemostat, the feeding was adapted MRS complemented with 7.5 g/L of maltose and 7.5 g/L of glucose. For AAMN, the feeding was adapted MRS complemented only with maltose (18.375 g/L) and regulation was performed by pulses of glucose, adding 0.15 g in the case of excess of LAB.

Finally, for the production proof of concept utilizing ECO* and SAC*, cultivations were performed at a stirring frequency of 1000 min, an air flow rate of 1 VVM with Verduyn medium at a pH of 6.8 and a dilution rate of 0.1 h^-1^. The population ratio for inoculation was 1:1 ECO*/SAC*. For the chemostat experimental design (uncontrolled) experiments, three different concentrations were used: 7.5/2.6 g/L GLC/XYL, 6.7/3.3 g/L GLC/XYL and 5.9/4.1 g/L GLC/XYL. For the AAMN experiment, two different runs were performed, and control rule was set to try to maintain the population around 50% by pulsing 0.25 g of either GLC or XYL every 15 minutes in the case of finding an excess of ECO*or SAC* respectively.

### FC Data treatments

Data treatment started by deleting the noise (zero and negative values) and doublets. To address doublet contamination, a linear regression analysis was conducted, correlating the area and height of the forward scattering signal (FSC-A and FSC-H). Data points with Pearson standardized residual values exceeding 2 were identified and eliminated. Samples exhibiting more than 5 % doublets or having a total remaining event count below 20,000 were excluded from further analysis. The fractions of events corresponding to each strain, their respective fluorescence and the time of for pump activation were calculated using the MiPI Flow Cytometry Analysis toolbox (mFCAtoolbox) part of the mBIOMAS core toolbox available at https://gitlab.uliege.be/mipi/published-software/mbiomas-core all installation and use information can be obtained there. Population fractions were based on gating the yeast populations, defined as events with higher Forward Scatter signal area (FSC-A) than 316227 (10^5.5^) where bacteria were defined as cells with lower values. All FC Data, and treatment algorithms and pipelines can be found in the repository https://gitlab.uliege.be/mipi/published-software/2024-cocultures

### Metabolite Analysis

Samples from fermentation experiments were processed for carbon sources (glucose, maltose, xylose and ethanol) measurement by HPLC with an Aminex HPX-87H column (Bio-Rad, Hercules CA, USA) at 45°C and 5 mM Sulfuric acid as mobile phase. An Agilent 1200 Series HPLC system was used with a refraction index detector (RID) at 50°C (Agilent, Santa Clara, CA, USA).

The L-tyrosine and p-CA were detected and quantified by UPLC measurements with Shimadzu Nexera series 40. The flow was 0.6 mL/min and the temperature was 40°C. Detection was done based on a RID and the column was a Water acquity C18 premier 1.7 μm, 2.1 x 50 mm. The analyses were operated with two mobile phases, 0.1 % trifluoroacetic acid (TFA) in methanol and 0.1 % TFA in water, the distribution table for the mobile phases for the HPLC analyses can be found in **Supplementary note 6**.

### Elementary Mode (EM) analysis

In this work different central carbon metabolism networks were constructed from genome and central core models and metabolic constraints available in the literature. Specifically, for ECO and ECO*, the models used were the ones published by Martinez *et al.*^71^, Edwards *et al.*^72^, Lendenmann *et al.*^73^, Covert *et al.*^74^, Peng *et al.*^75^, Price *et al.*^76^, Schmid *et al.*^77^ and Visser *et al.*^78^. For SAC and SAC*, we used the metabolic models developed by Rizzi *et al.*^79^, Carlson *et al.*^80^, Visser *et al.*^81^, Kesten *et al.*^82^, Lao-Martil *et al.*^83^ and Pitkanen *et al.*^84^. For KAZ, we used the metabolic models described in van Dijken *et al.*^85^, Middelhoven *et al.*^86^, Kurtzman *et al.*^87^, Balarezo-Cisneros *et al.*^88^, Guidot *et al.*^89^ and Dirick *et al.*^90^. Finally, for LAB, we used the metabolic models described in Wang *et al.*^91^, Tsuji *et al.*^92^, Poolman *et al.*^93^, Monedero *et al.*^94^, Koduru *et al.*^95^, Hickey *et al.*^96^, Hatti-Kaul *et al.*^97^, Filannino *et al.*^98^, Bai et *al.*^99^, Meng *et al.*^100^. Metabolic reactions for each central carbon metabolism were selected including transport reactions and a Biomass composition based mainly on the protein mean content and composition according to each strain bibliography (**Supplementary note 1**). With this selected and curated reactions a stoichiometric matrix was constructed for each organism and then separated into Elementary Flux Distributions with the aid of the program *efmtool* created by Terzer et al.^101^ and available at https://csb.ethz.ch/tools/software/efmtool.html. This program allowed us to construct the polyhedral cone of solutions for every strain containing all basal flux distributions in the cell. From these distributions we selected the most extremes for the metabolites that could be utilized by each strain. After that we performed and test comparison between EMs in the biomass/substrate yield (*Y_max_*) space for each of the EMs in which we compare the yields and derive the aggressivity or competitivity (*θ_i_*) of the strain for each substrate (*i*) between two different microorganisms (*a* and *b*) with the following formula:

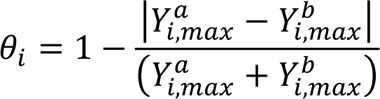

This formula effectively compares the capabilities of consumption for each substrate in equality of masses between the strains in a continuous culture where the growth rate is constrained to the dilution rate, it therefore, compares the maximum aggressivity of each strain to dominate the environment. If this index value is 1 it means that both microorganisms could transform the same amount of substrate by unit of biomass and therefore the equilibrium point in a continuous culture would be set by the amount of biomass in the system and their growth rates. Effectively changes in the concentration of this metabolite would not allow us to modify this point of equilibrium, or in other words this metabolite would not allow us to exert control in the system composition. Values less than 1 indicate an unbalanced competition in which one of the strains has a better efficiency of substrate conversion and therefore in continuous cultures with dilution rates smaller than the maximum growth rates then the strain with higher index would eventually dominate the environment, the higher is the disparity the faster this fate should occur. Finally, as this value approaches to 0 we arrive at a perfect exclusivity state in which only one of the strains can efficiently transform this substrate into biomass. Having alternative indexes for the microorganisms for two different substrates would be the perfect case for controllability as it would allow us to easily feed separately each biomass.

All used organisms curated reaction files and EMs analysis algorithms and pipelines can be found in the repository https://gitlab.uliege.be/mipi/published-software/2024-cocultures

### Dynamic metabolic flux modelling and analysis

To model the dynamics of the internal fluxes of the strains and their changes in metabolism, we used a hybrid cybernetic modelling approach. This approach has been thoroughly described in the literature and a detailed description of the approach used in this work can be found in the **Supplementary Note 3 and 4**. Briefly, this approach consists of time-differential equations for the metabolites consumption based on the rates of a selected set of EMs regulated by the relative enzymatic compromise of cells to each EMs given by a metabolic fitness objective, in this case growth rate. This approach allowed us to calculate the approximated fluxes through diverse metabolic processes such as balanced metabolite oxidations, metabolite fermentations and by-product consumptions.

The calculation of these models was performed with the MONCKS software published in a previous article^11^. In this work this software was translated to python and implemented through a more complete computational toolbox called MiPI Biomass Modelling and Simulation toolbox (mBIOMAS) which core is available at https://gitlab.uliege.be/mipi/published-software/mbiomas-core all installation and use information can be obtained there. The use of this software is simple and only requires the introduction of the EMs Yields and rates for each strain as well as the initial bioprocess conditions and in the case of the AAMN models the characteristics of the control rules and actuator.

In this work we mainly simulated ECO/SAC systems. First axenic cultures were performed to fit the parameters of the models concerning growth rates in the main substrates and by-product consumption, after that the chemostat and AAMN co-cultures of ECO/SAC were used to validate the models. Finally, simulations for the ECO/SAC system were performed by changing either the growth rates on each substrate (fitness) and by reducing the by-product formation (product yield reduction). The simulations by the explained perturbations allowed to study the effects of metabolic niche fitness and metabolic niche size on population stability and controllability. All the information regarding its use its available in the repository and all used files and simulation pipelines in this work are available in the repository https://gitlab.uliege.be/mipi/published-software/2024-cocultures.

To address competition and controllability between two different strains in this dynamic environment, we used the Shannon’s Entropy to measure the total distribution of metabolic niches across the different strains^102^. For these we use the cybernetic model data that allows us to calculate the volumetric consumption rates for all metabolites across the individual strains. By normalizing each metabolite consumption flow per each strain by the total consumption rate for each substrate s, we can approximate niche participation for each strain (Π*_s_^a^*). This niche participation can be set as the probability for a substrate pulse or environmental change would be processed into biomass by each one of the strains and their metabolic states. With this approximation we calculate the partial Entropy (h) of each strain (a and b) for a given substrate (s) in a by the following formulas:

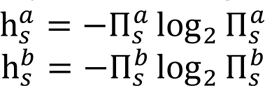

With this information, we can formulate a Monopoly index for each substrate (*M_s_*) by subtracting to 1 the sum of the partial entropies for all strains competing for the substrate:

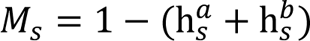

This index *M_s_* will be 0 when the probability of the s is the same for it to be consumed by either of the two strains 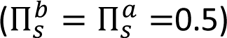and it tends to be 1 whenever the probability of the substrate is to be consumed by one of the strains is closer to 1.

Furthermore, we can calculate the total substrate competitivity for each substrate (Θ_*s*_) by comparing the partial entropies corresponding to each microorganism, and the systems total Entropy for each substrate (H_*s*_) which would be the summatory of the partial entropies of all microorganisms present in the system. This comparison of the difference between their entropies and the systems entropy by the competitivity index shows how well the substrates are being distributed along these microorganisms, and can be calculated as follows:

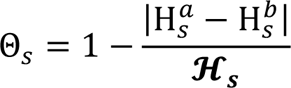

The competitivity of the system can be extended by the mean of all the individual competitivities and tends to a value of 0 for the cases when a substrate is being consumed by a single strain and tends to 1 when the substrate is being consumed in the exact same rate by both strains. In the latter case, from a control perspective there is no action possible around the substrate s (pulse or withdrawal) that would allow us to excerpt a meaningful change of the composition of the system. Conversely when this value tends to 1 then changes into this substrate would impact severely the composition of the systems biomass. Therefore, we defined the controllability of the system as K_*s*_ = 1 − Θ. The behaviour of these indexes can be better observed in the **Supplementary note 4.** The algorithms and pipelines used for the calculation of this indexes can be found in the repository https://gitlab.uliege.be/mipi/published-software/2024-cocultures

Finally, this entropy analysis was extended to all substrates or Metabolic Niches present in the environment and calculate the total Niche participations and to determine the total dynamic controllability of the system for the Chemostat and AAMN experiments and simulations.

## Data availability

The raw data sets are available on GitLab at https://gitlab.uliege.be/mipi/published-software/2024-cocultures.

Extra data, tables, graphs and documentation can be found in the **Supplementary Information** file separated as: **Supplementary Note #1**: Central Carbon Metabolic Network Models. **Supplementary Note #2**: Elementary Mode Analysis. **Supplementary Note #3**: MONCKS hybrid cybernetic modelling. **Supplementary Note #4**: Shannon entropy as an index for competitivity and controllability. **Supplementary Note #5**: Calculation of auto-aggregation. **Supplementary Note #6**: Media preparation and analysis specifications.

## Code availability

An updated version of MONCKS is provided on GitLab at https://gitlab.uliege.be/mipi/published-software/mbiomas-core.

## Acknowledgement

JAM is supported by a post-doctoral grant provided by the Service Public de Wallonie (SPW) and the H2020 program of the European commission (Era-Cobiotech project Contibio). RB is supported by a post-doctoral grant provided by the Service Public de Wallonie (SPW) for the Sunup project. TGSA is supported by a PhD grant provided by the “Fonds de la Recherche Scientifique” FRS-FNRS, from the Walloon region of Belgium. FD received funding from a research grant provided by the Service Public de Wallonie (SPW) and the H2020 program of the European commission (Era-Cobiotech project Contibio). The participation of Caheri Salas in the construction of the yeast p-coumaric acid production strain is acknowledged.

## Author contributions

JAM performed the experiments related to ECO-SAC co-culture, developed the computational toolbox, and drafted the manuscript. RB performed the experiments related to LAB-KAZ and ECO*-SAC*. TGSA performed the experiments related ECO*-SAC* to produce p-coumaric acid. GG designed the strains involved in ECO*-SAC*. LMM and LC constructed the ECO*-SAC* strains. FD captured grants and funding, developed the concept of metabolic niches, supervised the research and wrote the manuscript.

## Competing interests

The authors declare no competing interests.

## Notes

### Competing Interest Statement

The authors have declared no competing interest.

