## Supplementary material for "Automated adjustment of metabolic niches enables the control of natural and engineered microbial co-cultures"

In this file all supplementary information relative to the previously mentioned manuscript is to be found accordingly to the following **index**:

- **PAGE 02 -Supplementary Note #1**: Central Carbon Metabolic Network Models.
- **PAGE 10 -Supplementary Note #2**: Elementary Mode Analysis.
- **PAGE 19 -Supplementary Note #3**: MONCKS hybrid cybernetic modelling.
- **PAGE 29 -Supplementary Note #4**: Shannon entropy as an index for competitivity and controllability.
- **PAGE 31 -Supplementary Note #5**: Calculation of auto-aggregation.
- **PAGE 34 -Supplementary Note #6**: Media preparation and analysis specifications.

### Supplementary Note #1: Central Carbon Metabolic Network Models

In this note, we present the central carbon metabolism networks constructed from genome and central core models and metabolic constraints available in the literature. Specifically for ECO and ECO* the models used were the ones published by Martinez *et al.* [1], Edwards *et al.* [2], Lendenmann *et al.* [3], Covert *et al.* [4], Peng *et al.* [5], Price *et al.* [6], Schmid *et al.* [7], Visser *et al.* [8]. For SAC and SAC* Rizzy *et al.* [9], Carlson *et al.* [10], Viser *et al.* [11], Kesten *et al.* [12], Martil *et al.* [13] and Pitkanen [14]. For KAZ, van Dijken *et al.* [15], Middelhoven *et al.* [16], Kurtzman *et al.* [17], Balarezo-Cisneros *et al.* [18], Guidot *et al.* [19] and Dirick *et al.* [20]. Finally, for LAB Wang *et al.* [21], Tsuji *et al.* [22], Poolman *et al.* [23], Monedero *et al.* [24], Koduru *et al.* [25], Hickey *et al.* [26], Hatti-Kaul *et al.* [27], Filannino *et al.* [28], Bai *et al.* [29], Meng *et al.* [30].

Lists of reactions used in this work for ***Escherichia coli* W3110 (ECO)**

**# GLYCOLYSIS**
*RGLCx :* GLCx + PEP = G6P + PYR
*RGLCx2 :* GLCx + ATP = G6P
*R2r :* G6P = F6P
*R3a :* F6P + ATP = FBP
*R3b :* FBP = F6P
*R4r :* FBP = GAP + DHAP
*R5r :* GAP = DHAP
*R6r :* GAP = BPG + NADH + ATP
*R7r :* BPG = PEP
*R8a :* PEP = ATP + PYR
*R8b :* 2 ATP + PYR = PEP
**# GLYCEROL METABOLISM**
*R9 :* DHAP + NADH = G3P
*R10 :* G3P = DHAP + QH2
*R11r :* GLY = G3P
*RGLYx :* GLYx = GLY
**# PENTOSE PHOSPHATE PATHWAYS**
*R12 :* G6P = GAP + PYR + NADPH
*R13 :* G6P = RU5P + 2 NADPH + CO2
*R14r :* RU5P = X5P
*R15r :* RU5P = R5P
*R16r :* R5P + X5P = S7P + GAP
*R17r :* X5P + E4P = F6P + GAP
*R18r :* S7P + GAP = F6P + E4P
**# PYRUVATE METABOLISM**
*R19 :* PYR = ACCOA + NADH + CO2
*R20r :* ACCOA = ACTP
*R21r :* ACTP = ACE + ATP
*R22r :* ACCOA + 2 NADH = ETH
*RETHx :* ETHx = ETH
*R23 :* PYR = ACE + QH2
*R25 :* 2 ATP + ACE = ACCOA
*RACEx :* ACEx = ACE
**# TRICARBOXYLIC ACID CYCLE**
*R37 :* OAA + ACCOA = CIT
*R38r :* CIT = ICIT
*R39r :* ICIT = AKG + NADPH + CO2
*R40a :* AKG = SUCCOA + NADH + CO2
*R40br :* SUCCOA = SUC + ATP
*R41r :* SUC = FUM + QH2
*R42r :* FUM = MAL
*R43r :* MAL = OAA + NADH
*R45 :* NADPH = NADH
*R46 :* QH2 = 2 HEXT
*R47 :* NADH = 2 HEXT + QH2
*R48 :* 1 HEXT + NADH = NADPH
*R49 :* 3 HEXT = ATP
**# ANAPLEROTIC REACTIONS**
*RSUCx :* SUCx = SUC
*R55 :* OAA + ATP = PEP + CO2
*R56 :* PEP + CO2 = OAA
*R57 :* ICIT = GLX + SUC
*R58 :* ACCOA + GLX = MAL
*R59a :* MAL = PYR + NADH + CO2
*R59b :* MAL = PYR + NADPH + CO2
**# MAINTENANCE**
*RMNT :* ATP = MAINT
**# BIOMASS FORMATION**
*BIOMSynth* : 0.06156 G6P + 0.70798 R5P + 0.47711 E4P + 2.4010 PEP + 3.647613 PYR + 1.108136 ACCOA + 1.323606 AKG + 2.139317 OAA + 44.95644 ATP + 13.1745 NADH = BIOM + 0.538677 CO2

***NOTE:** reaction names ending with r indicate reversible reactions; the ones ending with x indicate transport reactions. Metabolite names with an x at the end indicate external metabolites, all else are cytosolic metabolites.

Lists of reactions used in this work for ***Saccharomyces cerevisiae*** **CENPK** ***pchi-eGFP*** **(SAC).**

**# GLYCOLYSIS**
*RGLCx :* GLCx + ATP = G6P
*R2r :* G6P = F6P
*R3r :* F6P + ATP = FBP
*R4r :* FBP = GAP + DHAP
*R5r :* GAP = DHAP
*R6r :* GAP = BPG + NADH + ATP.
*R7r :* BPG = PEP
*R8r :* PEP = PYR + ATP

**# GLYCEROL METABOLISM**
*R9 :* DHAP + NADH = G3P
*R10 :* G3P = DHAP + 1.2 ATPm
*R11r :* G3P = GOL
*RGOLx :* GOLx = GOL

**# PENTOSE PHOSPHATE PATHWAYS**
*R13 :* G6P = RU5P + 2 NADPH + CO2
*R14r :* RU5P = X5P
*R15r :* RU5P = R5P
*R16r :* R5P + X5P = S7P + GAP
*R17r*: X5P + E4P = F6P + GAP
*R18r :* S7P + GAP = F6P + E4P

**# PYRUVATE METABOLISM**
*R19 :* PYR = ACD + CO2
*R20 :* PYR + ATP + CO2 = OAA
*R21 :* ACD + NADH = ETH
*R22 :* ACD + NADHm = ETH
*RETHx :* ETH = ETHx
*R23r :* ACD = ACE + NADPH
*RACEx*: ACE = ACEx
*R25r :* ACE + 2 ATP = ACCOA

**# MITOCHONDRIAL TRANSPORT**
*R26 :* ATPm = ATP
*R27 :* PYR = PYRm
*R28r*: ACCOA = ACCOAm
*R29 :* MALm = MAL
*R30 :* FUMm + SUC = FUM + SUCm
*R31 :* CITm + MAL = CIT + MALm
*R32 :* SUC = SUCm
*R33 :* AKG + MALm = AKGm + MAL
*R34r :* OAA + NADH = OAAm + NADHm.
*R35r :* ETH = ETHm

**# MITOCHONDRIAL METABOLISM**
*R36 :* PYRm = ACCOAm + NADHm + CO2
*R37 :* OAAm + ACCOAm = CITm
*R38r :* CITm = ICITm
*R39ar :* ICITm = AKGm + NADHm + CO2
*R39br :* ICITm = AKGm + NADPHm + CO2
*R40 :* AKGm = SUCm + NADHm + ATPm + CO2
*R41r :* SUCm = FUMm + 0.5 NADHm
*R42r :* FUMm = MALm
*R43r :* MALm = OAAm + NADHm
*R44 :* MALm = PYRm + NADHm
*R45 :* NADHm = ATPm
*R46 :* ETHm + 2 ATPm = ACCOAm + 2 NADHm

**# CYTOSOL METABOLISM**
*R47 :* OAA + ACCOA = CIT
*R48r :* CIT = ICIT
*R49 :* ICIT = AKG + NADPH + CO2
*R50 :* ICIT = GLX + SUC
*R51 :* GLX + ACCOA = MAL
*R52r :* OAA + NADH = MAL
*R53r :* FUM + 0.5 NADH = SUC
*R54r :* FUM = MAL
*R55 :* OAA + ATP = PEP + CO2
*RSUCx :* SUC = SUCx
*R57 :* CIT + ATP = ACCOA + OAA
*R58 :* OAA + ATP = PEP + CO2

**#MAINTEINANCE**
*RMNT :* ATP = MAINT

**# BIOMASS FORMATION**
*RBIO :* 1.04 AKG + 0.57 E4P + 0.11 GOL + 2.39 G6P + 1.07 OAA + 0.99 PEP + 0.57 BPG + 1.15 PYR + 0.74 R5P + 2.36 ACCOA + 0.31 ACCOAm + 11.55 NADPH + 1.51 NADPHm + 30.48 ATP + 0.43 CO2 = BIOM + 2.68 NADH + 0.53 NADHm

***NOTE:** reaction names ending with r indicate reversible reactions; ending with x indicate transport reactions. Metabolite names with an x at the end indicate external metabolites, all else are cytosolic metabolites.

Lists of reactions used in this work for ***Lactobacillus plantarum*** **(LAB).**

**# Transport reactions**

*RMLTr :* MLTx = MLT
*RGLC :* GLCx + ATP = G6P
*RGOL :* GOLx = GOL
*RETHr :* ETH = ETHx
*RACEr :* ACE = ACEx
*RLACr :* LAC = LACx
*RALAr :* ALA = ALAx
*RARG :* ARGx = ARG
*RASN :* ASNx = ASN
*RASP :* ASPx = ASP
*RCYS :* CYSx = CYS
*RGLN :* GLNx = GLN
*RGLUr :* GLU = GLUx
*RGLYr :* GLY = GLYx
*RHIS :* HISx = HIS
*RILEr :* ILEx = ILE
*RLEUr :* LEUx = LEU
*RLYS :* LYSx = LYS
*RMET :* METx = MET
*RPHEr :* PHEx = PHE
*RPRO :* PROx = PRO
*RSERr :* SERx = SER
*RSUCr :* SUCx = SUC
*RTHR :* THRx = THR
*RTRP :* TRPx = TRP
*RTYR :* TYRx = TYR
*RVALr :* VALx = VAL

**# GLYCOLYSIS**

*R2r :* G6P = F6P
*R3 :* F6P + ATP = 2 GAP
*R4r :* GAP = PG3 + NADH + ATP
*R5r :* PG3 = PEP
*R6r :* PEP = PYR + ATP

**# PENTOSE PHOSPHATE PATHWAY**

*R7 :* G6P = RU5P + CO2 + 2 NADPH
*R8r :* RU5P = XU5P
*R9r :* RU5P = R5P
*R10r :* R5P + XU5P = S7P + GAP
*R11r :* S7P + GAP = F6P + E4P
*R12r :* XU5P + E4P = F6P + GAP

**# PYRUVATE and FERMENTATION**

*R13 :* PYR + ATP + CO2 = OAA
*R14 :* PYR = ACCOA + CO2 + NADH
*R15 :* PYR = ACD + CO2
*R16 :* PYR + NADH = LAC
*R18 :* ACD + NADH = ETH
*R19 :* ACD = ACE + NADPH
*R20 :* ACE + 2 ATP = ACCOA

**# CITRIC ACID CYCLE**

*R21 :* OAA + ACCOA = CIT
*R22r :* CIT = ICIT
*R23 :* ICIT = AKG + CO2 + NADPH
*R24 :* AKG = SUC + NADH + CO2
*R25r :* SUC = MAL + 0.5 NADH

*R28 :* MAL = LAC + CO2
*R29 :* MAL = PYR + CO2 + NADH

**# MALTOSE, XYLOSE and GYCEROL**

*R30 :* MLT = GLCx + G6P
*R31 :* MLT = 2 GLCx
*R35 :* GOL + ATP = PG3

**## BIOMASS COMPONENTS**

**# AMINO ACIDS**

*R37 :* PYR + NADPH = ALA
*R38 :* AKG + 4 NADPH + 7 ATP = ARG + NADH
*R39 :* OAA + NADPH + 3 ATP = ASN
*R40 :* OAA + NADPH = ASP
*R41 :* PG3 + 4 NADPH + 7 ATP = CYS + NADH
*R42 :* AKG + NADPH = GLU
*R43 :* AKG + NADPH + ATP = GLN
*R45 :* R5P + 4 NADPH + 6 ATP + CO2 = HIS + 3 NADH
*R48 :* ACCOA + AKG + 4 NADPH + 2 ATP = LYS + 2 NADH
*R49 :* OAA + 11 NADPH + 7 ATP + CO2 = MET
*R51 :* AKG + 3 NADPH + ATP = PRO
*R53 :* OAA + 3 NADPH + 2 ATP = THR
*R54 :* PEP + E4P + PG3 + 3 NADPH + 5 ATP = TRP + 2 NADH
*R55 :* 2 PEP + E4P + 2 NADPH + ATP = TYR + NADH
*R57 :* 0.086 ALA + 0.041 ARG + 0.031 ASP + 0.059 ASN + 0.034 CYS + 0.036 GLU + 0.064 GLN + 0.092 GLY + 0.015 HIS + 0.061 ILE + 0.087 LEU + 0.072 LYS + 0.025 MET + 0.038 PHE + 0.035 PRO + 0.051 SER + 0.056 THR + 0.017 TRP + 0.027 TYR + 0.072 VAL + 4.306 ATP = PROT

**# ENERGY EXCHANGES**

*RMNT :* ATP = MAINT

**# BIOMASS**

*RBIO :* 4.2 PROT = BIOM

***NOTE:** reaction names ending with r indicate reversible reactions; ending with x indicate transport reactions. Metabolite names with an x at the end indicate external metabolites, all else are cytosolic metabolites.

Lists of reactions used in this work for ***Kazachstania bulderi*** **(KAZ).**

**# Transport reactions**

*RGLC :* GLCx + ATP = G6P
*RGOL :* GOL = GOLx
*RETHr :* ETH = ETHx
*RACEr :* ACE = ACEx
*RSUCr :* SUC = SUCx
*RALAr :* ALAx = ALA
*RARG :* ARGx = ARG
*RASN :* ASNx = ASN
*RASP :* ASPx = ASP
*RCYS :* CYSx = CYS
*RGLN :* GLNx = GLN
*RGLUr :* GLUx = GLU
*RGLYr :* GLYx = GLY
*RHIS :* HISx = HIS
RILEr : ILEx = ILE
*RLEUr :* LEUx = LEU
*RLYS :* LYSx = LYS
*RMET :* METx = MET
*RPHEr :* PHEx = PHE
*RPRO :* PROx = PRO
*RSERr : SERx = SER*
*RTHR :* THRx = THR
*RTRP :* TRPx = TRP
*RTYR :* TYRx = TYR
*RVALr :* VALx = VAL

**# GLYCOLYSIS**

*R2r :* G6P = F6P
*R3 :* F6P + ATP = 2 GAP
*R4r :* GAP = PG3 + NADH + ATP
*R5r :* PG3 = PEP
*R6r :* PEP = PYR + ATP

**# PENTOSE PHOSPHATE PATHWAY**

*R7 :* G6P = RU5P + CO2 + 2 NADPH
*R8r*: RU5P = XU5P
*R9r :* RU5P = R5P
*R10r*: R5P + XU5P = S7P + GAP
*R11r*: S7P + GAP = F6P + E4P
*R12r*: XU5P + E4P = F6P + GAP

**# PYRUVATE and FERMENTATION**

*R13*: PYR + ATP + CO2 = OAA
*R14*: PYR = ACCOA + CO2 + NADH
*R15*: PYR = ACD + CO2
*R18*: ACD + NADH = ETH
*R19*: ACD = ACE + NADPH
*R20 :* ACE + 2 ATP = ACCOA

**# CITRIC ACID CYCLE**

*R21 :* OAA + ACCOA = CIT
*R22r :* CIT = ICIT
*R23 :* ICIT = AKG + CO2 + NADPH
*R24 :* AKG = SUC + ATP + CO2 + NADH
*R25r :* SUC = MAL + 0.5 NADH
*R26r :* MAL = OAA + NADH

**# XYLOSE and GLYCEROL**

*R34 :* GAP + NADH = GOL

**b**

*R37 :* PYR + NADPH = ALA
*R38 :* AKG + 4 NADPH + 7 ATP = ARG + NADH
*R39 :* OAA + NADPH + 3 ATP = ASN
*R40 :* OAA + NADPH = ASP
*R41 :* PG3 + 4 NADPH + 7 ATP = CYS + NADH
*R42 :* AKG + NADPH = GLU
*R43 :* AKG + NADPH + ATP = GLN
*R44 :* PG3 = GLY + NADH + 2 NADPH + CO2
*R45 :* R5P + 4 NADPH + 6 ATP + CO2 = HIS + 3 NADH
*R46 :* OAA + PYR + 5 NADPH + 2 ATP = ILE
*R47 :* ACCOA + 2 PYR + 2 NADPH = LEU + *NADH*
*R48 :* ACCOA + AKG + 4 NADPH + 2 ATP = LYS + 2 NADH
*R49 :* OAA + 11 NADPH + 7 ATP + CO2 = MET
*R50 :* 2 PEP + E4P + 2 NADPH + ATP = PHE
*R51* : AKG + 3 NADPH + ATP = PRO
*R52* : PG3 + NADPH = SER + NADH
*R53* : OAA + 3 NADPH + 2 ATP = THR
*R54 :* PEP + E4P + PG3 + 3 NADPH + 5 ATP = TRP + 2 NADH
*R55 :* 2 PEP + E4P + 2 NADPH + ATP = TYR + NADH
*R56 :* 2 PYR + NADPH = VAL
*R57 :* 0.047 ALA + 0.056 ARG + 0.025 ASN + 0.102 ASP + 0.009 CYS + 0.107 GLU + 0.027 GLN + 0.056 GLY + 0.091 HIS + 0.052 ILE + 0.079 LEU + 0.031 LYS + 0.047 MET + 0.04 PHE + 0.043 PRO + 0.052 SER + 0.014 THR + 0.04 TRP + 0.065 TYR + 0.017 VAL + 4 ATP = 4.825 PROT.

**# OXIDATIVE PHOSPHORYLATION and ENERGY EXCHANGES**

RMNT : ATP = MAINT

**# BIOMASS**

RBIO : 0.47 PROT + 0.003 GOL = BIOM

***NOTE:** reaction names ending with r indicate reversible reactions; ending with x indicate transport reactions. Metabolite names with an x at the end indicate external metabolites, all else are cytosolic metabolites.

Lists of reactions used in this work for ***Escherichia coli* WHIC *+pJLBaroGfbr +pTrcTyrCpheACM* (ECO*)**

**# GLYCOLYSIS**
*RGLCx2* : GLCx + ATP = G6P
*RXYL :* XYLx + HEXT = XYL
*R2r :* G6P = F6P
*R3a :* F6P + ATP = FBP
*R3b :* FBP = F6P
*R4r :* FBP = GAP + DHAP
*R5r :* GAP = DHAP
*R6r :* GAP = BPG + NADH + ATP
*R7r :* BPG = PEP
*R8a :* PEP = ATP + PYR
*R8b :* 2 ATP + PYR = PEP
**# GLYCEROL METABOLISM**
*R9 :* DHAP + NADH = G3P
*R10 :* G3P = DHAP + QH2
*R11r :* GOL = G3P
*RGOLx :* GOLx = GOL
**# PENTOSE PHOSPHATE PATHWAYS**
*R12 :* G6P = GAP + PYR + NADPH
*R13 :* G6P = RU5P + 2 NADPH + CO2
*R14r :* RU5P = X5P
*R14b :* XYL + ATP = X5P + NADPH
*R15r :* RU5P = R5P
*R16r :* R5P + X5P = S7P + GAP
*R17r :* X5P + E4P = F6P + GAP
*R18r :* S7P + GAP = F6P + E4P
**# PYRUVATE METABOLISM**
*R19 :* PYR = ACCOA + NADH + CO2
*R20r :* ACCOA = ACTP
*R21r :* ACTP = ACE + ATP
*R22r :* ACCOA + 2 NADH = ETH
*RETHx :* ETHx = ETH
*R23 :* PYR = ACE + QH2
*R25 :* 2 ATP + ACE = ACCOA
*RACEx :* ACEx = ACE
**# TRICARBOXYLIC ACID CYCLE**
*R37 :* OAA + ACCOA = CIT
*R38r :* CIT = ICIT
*R39r :* ICIT = AKG + NADPH + CO2
*R40a :* AKG = SUCCOA + NADH + CO2
*R40br :* SUCCOA = SUC + ATP
*R41r :* SUC = FUM + QH2
*R42r :* FUM = MAL
*R43r :* MAL = OAA + NADH
*R45 :* NADPH = NADH
*R46 :* QH2 = 2 HEXT
*R47 :* NADH = 2 HEXT + QH2
*R48 :* 1 HEXT + NADH = NADPH
*R49 :* 3 HEXT = ATP
**# ANAPLEROTIC REACTIONS**
*RSUCx* : SUCx = SUC
*R55 :* OAA + ATP = PEP + CO2
*R56 :* PEP + CO2 = OAA
*R57 :* ICIT = GLX + SUC
*R58 :* ACCOA + GLX = MAL
*R59a :* MAL = PYR + NADH + CO2
*R59b :* MAL = PYR + NADPH + CO2
**# Production Plasmids**
*R60 :* E4P + 2 PEP + 2 NADPH + ATP = TYRx
**# MAINTENANCE**
*RMNT :* ATP = MAINT
**# BIOMASS FORMATION**
*BIOMSynth :* 0.06156 G6P + 0.70798 R5P + 0.47711 E4P + 2.4010 PEP + 3.647613 PYR + 1.108136 ACCOA + 1.323606 AKG + 2.139317 OAA + 44.95644 ATP + 13.1745 NADH = BIOM + 0.538677 CO2

***NOTE:** reaction names ending with r indicate reversible reactions; ending with x indicate transport reactions. Metabolite names with an x at the end indicate external metabolites, all else are cytosolic metabolites.

Lists of reactions used in this work for ***Saccharomyces cerevisiae*** **PTA 408 +*pCA17* (SAC*)**

**# GLYCOLYSIS**
*RGLCx :* GLCx + ATP = G6P .
*R2r :* G6P = F6P .
*R3r :* F6P + ATP = FBP .
*R4r :* FBP = GAP + DHAP .
*R5r*: GAP = DHAP .
*R6r :* GAP = BPG + NADH + ATP.
*R7r :* BPG = PEP .
*R8r :* PEP = PYR + ATP .
**# GLYCEROL METABOLISM**
*R9 :* DHAP + NADH = G3P .
*R10 :* G3P = DHAP + 1.2 ATPm .
*R11r :* G3P = GOL .
*RGOLx :* GOLx = GOL .
**# PENTOSE PHOSPHATE PATHWAYS**
*R13 :* G6P = RU5P + 2 NADPH + CO2 .
*R14r :* RU5P = X5P .
*R15r :* RU5P = R5P .
*R16r :* R5P + X5P = S7P + GAP .
*R17r :* X5P + E4P = F6P + GAP .
*R18r :* S7P + GAP = F6P + E4P .
**# PYRUVATE METABOLISM**
*R19 :* PYR = ACD + CO2 .
*R20 :* PYR + ATP + CO2 = OAA .
*R21 :* ACD + NADH = ETH .
*R22*: ACD + NADHm = ETH .
*RETHx :* ETH = ETHx .
*R23r :* ACD = ACE + NADPH .
*RACEx : ACE = ACEx .*
*R25r*: ACE + 2 ATP = ACCOA .
**# MITOCHONDRIAL TRANSPORT**
*R26 :* ATPm = ATP .
*R27 :* PYR = PYRm .
*R28r*: ACCOA = ACCOAm .
*R29 :* MALm = MAL .
*R30*: FUMm + SUC = FUM + SUCm .
*R31 :* CITm + MAL = CIT + MALm .
*R32 :* SUC = SUCm .
*R33 :* AKG + MALm = AKGm + MAL .
*R34r :* OAA + NADH = OAAm + NADHm.
*R35r :* ETH = ETHm .
**# MITOCHONDRIAL METABOLISM**
*R36 :* PYRm = ACCOAm + NADHm + CO2 .
*R37 :* OAAm + ACCOAm = CITm .
*R38r :* CITm = ICITm .
*R39ar :* ICITm = AKGm + NADHm + CO2 .
*R39br*: ICITm = AKGm + NADPHm + CO2 .
*R40 :* AKGm = SUCm + NADHm + ATPm + CO2
. *R41r :* SUCm = FUMm + 0.5 NADHm .
*R42r :* FUMm = MALm .
*R43r :* MALm = OAAm + NADHm .
*R44 :* MALm = PYRm + NADHm .
*R45 :* NADHm = ATPm .
*R46 :* ETHm + 2 ATPm = ACCOAm + 2 NADHm .

**# CYTOSOL METABOLISM**
*R47 :* OAA + ACCOA = CIT .
*R48r :* CIT = ICIT .
*R49 :* ICIT = AKG + NADPH + CO2 .
*R50 :* ICIT = GLX + SUC .
*R51 :* GLX + ACCOA = MAL .
*R52r :* OAA + NADH = MAL .
*R53r :* FUM + 0.5 NADH = SUC .
*R54r :* FUM = MAL .
*R55 :* OAA + ATP = PEP + CO2 .
*RSUCx :* SUC = SUCx .
*R57 :* CIT + ATP = ACCOA + OAA .
*R58 :* OAA + ATP = PEP + CO2 .
**# PLASMID GENES**
*RTYR :* TYRx = TYR .
*RCMA :* TYR = CMAx .
**#MAINTEINANCE**
*RMNT*: ATP = MAINT .
**# BIOMASS FORMATION**

RBIO : 1.04 AKG + 0.57 E4P + 0.11 GOL + 2.39 G6P + 1.07 OAA + 0.99 PEP + 0.57 BPG + 1.15 PYR + 0.74 R5P + 2.36 ACCOA + 0.31 ACCOAm + 11.55 NADPH + 1.51 NADPHm + 30.48 ATP + 0.43 CO2 = BIOM + 2.68 NADH + 0.53 NADHm .

***NOTE:** reaction names ending with r indicate reversible reactions; ending with x indicate transport reactions. Metabolite names with an x at the end indicate external metabolites, all else are cytosolic metabolites.

#### *Supplementary Note 1 references*

27. Hatti-Kaul, R., Chen, L., Dishisha, T. & Enshasy, H. E. Lactic acid bacteria: from starter cultures to producers of chemicals. FEMS Microbiol Lett 365, (2018).

28. Filannino, P. et al. Metabolic responses of Lactobacillus plantarum strains during fermentation and storage of vegetable and fruit juices. Appl Environ Microbiol 80, 2206–2215 (2014).

29. Bai, D.-M., Zhao, X.-M., Li, X.-G. & Xu, S.-M. Strain improvement and metabolic flux analysis in the wild-type and a mutant Lactobacillus lactis strain for L(+)-lactic acid production. Biotechnol Bioeng 88, 681–689 (2004).

30. Meng, L. et al. The nutrient requirements of Lactobacillus acidophilus LA-5 and their application to fermented milk. J Dairy Sci 104, 138–150 (2021).

#### Supplementary Note #2: Elementary Mode Analysis.

The Elementary Mode (EM) analysis is based on the deconvolution of an organism's metabolic network into flux distribution networks that cannot be further reduced or simplified and that tie the consumption and productions of metabolites. Each one of these unique irreducible flux distributions is called an Elementary Mode (EM), and the collection of all EMs then represents all the organism's metabolic possibilities in the case of a full genomic scale Metabolic Network. In this work, however, we decided to utilize core central carbon metabolism Network simplifications instead of genome scale, since in all cases would be working in minimal media with few defined substrates. Even with just the core carbon reactions EMs number can reach values of several millions depending mostly on connectivity and reversibility of reactions extending to genome scale models in full then requires exhaustive computational power and time, and in our case probably rendering few or little more information given the constrains of our experiments, minimal media, few substrates, and continuous culture selecting certain growth rate requirements. Indeed, the production of secondary metabolites could be relevant in terms of the production of toxins against other organisms or certain types of quorum sensing compounds, between others, however the flux distributions towards these compounds is always expected to be small, affecting the main central carbon fluxes marginally, and furthermore, given the continuous culture conditions we expect their concentrations to be near 0 reducing their toxic or cellular effects greatly. Furthermore, having genomic scales hybrid cybernetic models is until now virtually impossible given the extensive computational power it would require, and in this case probably giving little more information to our case study than a well curated simplified core Metabolic Network that allowed us to understand and measure the effects of substrate temporality in the environment towards organism's niche specialization through phenotypic diversification.

In the **Supplementary note 1,** all used metabolic networks for all microorganisms used can be obtained. From their reaction description all EMs for each microorganism were obtained with the use of the computational tool *efmtools* developed by Terzer (2009) [1], available at https://csb.ethz.ch/tools/software/efmtool.html. We used for this work the MATLAB version through scripts designed as follows:

```
disp('--------------------------------')
disp('Calculating elementary modes by efmtool')
disp('--------------------------------')
   file = "./MetabolicNetwork.txt"
   ex = parse(file);
   mnet.stoich=ex.st;
   mnet.reversibilities = 1-ex.irrev_react;
   currdir = pwd;
   efmtool.path="C:\path to downloaded toolbox\efmtool";
   cd(efmtool.path);
   mnet=CalculateFluxModes(mnet);
   ex.ems=mnet.efms;
   cd(currdir);

   EFMmodel.m = ex.int_met; % names of intracellular metabolites
   EFMmodel.x = ex.ext_met; % names of extracellular metabolites
   EFMmodel.r = ex.react_name; % reaction names
```

```
   EFMmodel.sm = ex.st; % stoichiometric matrix (rows correspond to internal metabolites, columns to reactions)
   EFMmodel.sx = ex.ext; % same structure as st, but rows correspond to external metabolites
   EFMmodel.z = ex.ems; % elementary modes
   EFMmodel.sxz = EFMmodel.sx*EFMmodel.z;
save("EFMmodel.mat","EFMmodel");
```

From the operation of these MATLAB algorithms all EMs were obtained and then analysed by a subsequent python code. First this code reads the EMs, and then reduce their number by a set of simple constraints that depended on the microorganism pair and the culture conditions. For ECO/SAC and ECO*/SAC* pairs two constraints were used, growing EMs and Aerobic EMs, this is achieved by selecting EMs with BIOx reactions greater than 0 and with Oxygen evolution greater than 0. For the case of LAB/KAZ the constraints were growing and anaerobic, however, no aerobic EMs were found as expected given their genome and metabolic capabilities.  After this number reduction the system calculates the yields for each external metabolite in terms of biomass and selects the greater consumption ones for the calculation of their aggressivity and their competitivity (θ) as presented in the **Methods** section.  Finally, the program selects the simplified active set from the different Yields spaces, based on the experimental observations regarding the substrates supplemented to the growth media and all mayorly found by products and their consumption capabilities. For ECO and SAC the active set was selected containing: The highest biomass yield on GLC; the highest ETH production yield on GLC; the highest ACE production yield on GLC; the highest biomass yield on ETH and the highest biomass yield on ACE. For ECO* and SAC* the active set was the same as the latter ECO/SAC sets but with the addition of the highest biomass yield on XYL; the highest ETH production yield on XYL; the highest ACE production yield on XYL. None of the later resulted on an EMs for SAC* due to its impossibility to internalize XYL. LAB and KAZ were not simulated in this work given that we didn't have enough concentration information of amino acid contents in media to select the appropriate set. Finally, the algorithm calculates and outs different .csv files containing the selected sets and all EMs Yield spaces, along with some figures summarizing the model information. All algorithms, files and outs can be found in the Gitlab repository at https://gitlab.uliege.be/mipi/published-software/2024-cocultures.  In this Note we will introduce the main resulting figures for each microorganism pair.

***Supplementary Note 2 references:***


1. Terzer, Marco. Large scale methods to enumerate extreme rays and elementary modes. ETH, Doctoral Thesis. Diss., Eidgenössische Technische Hochschule ETH Zürich, Nr. 18538, (2009). https://doi.org/10.3929/ethz-a-005945733

#### *Supplementary Note 2 Figures and Tables:*

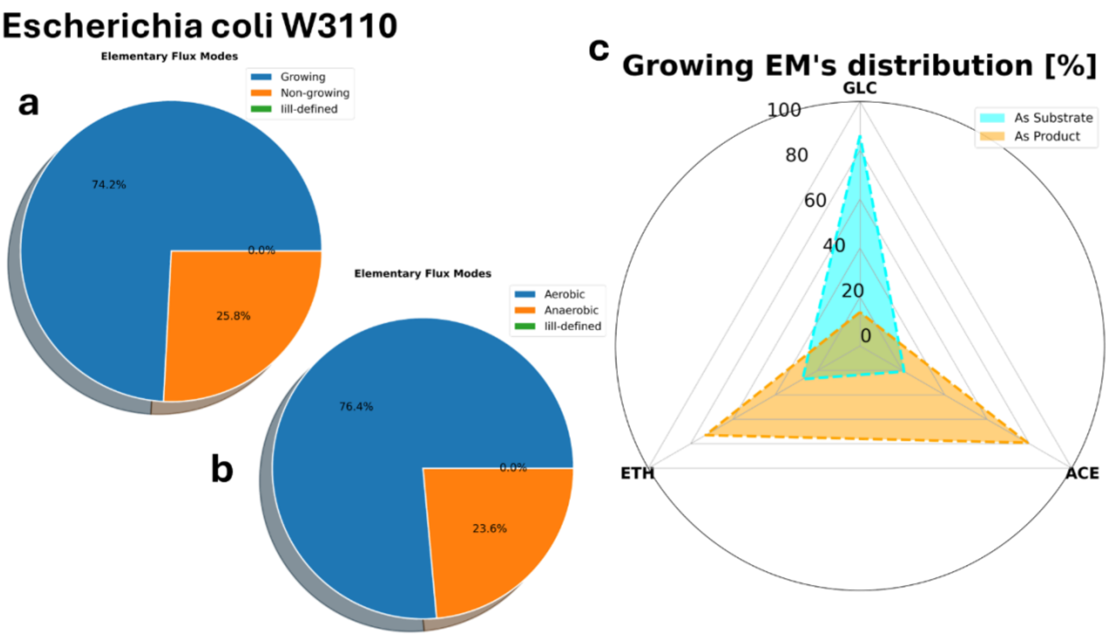


**Supplementary Figure 1** a) Distribution of EMs under the growth constraint for strain ECO. b) Distribution of EMs under the constraint of Aerobiosis. c) Distribution of EMs under the constraints of single substrate consumption and principal produced compounds.

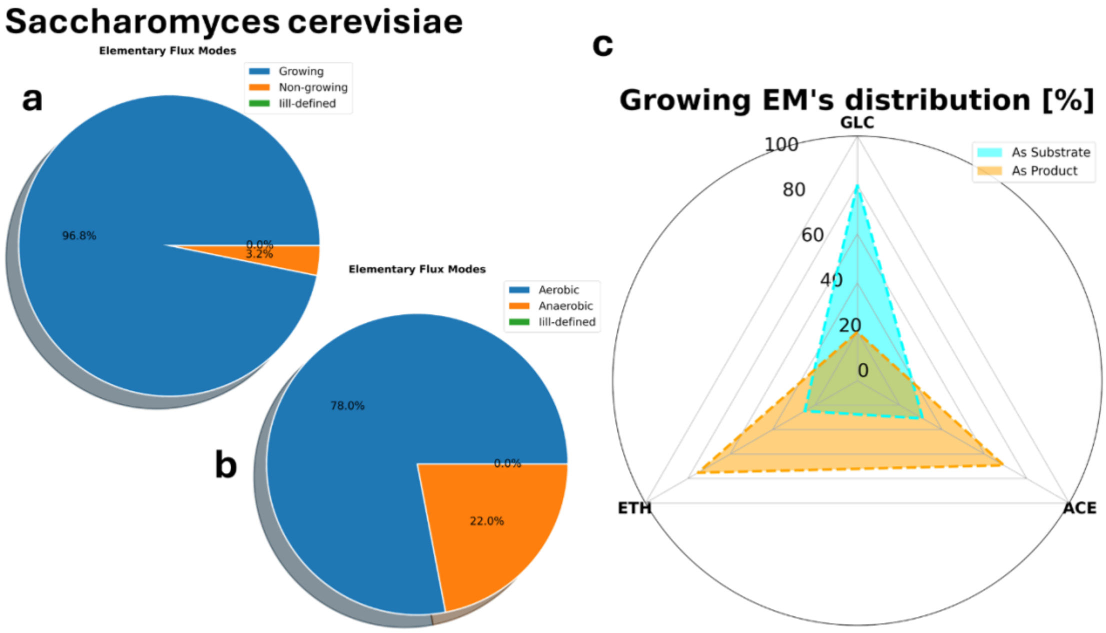


**Supplementary Figure 2** a) Distribution of EMs under the growth constraint for strain SAC. b) Distribution of EMs under the constraint of Aerobiosis. c) Distribution of EMs under the constraints of single substrate consumption and principal produced compounds.

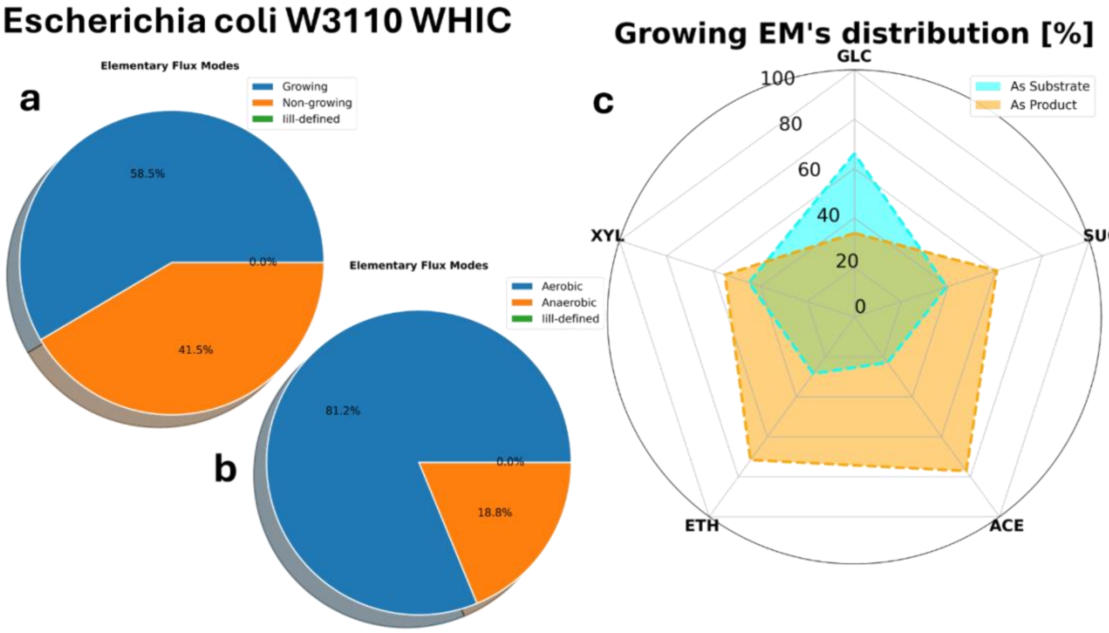


**Supplementary Figure 3** a) Distribution of EMs under the growth constraint for strain ECO*. b) Distribution of EMs under the constraint of Aerobiosis. c) Distribution of EMs under the constraints of single substrate consumption and principal produced compounds.

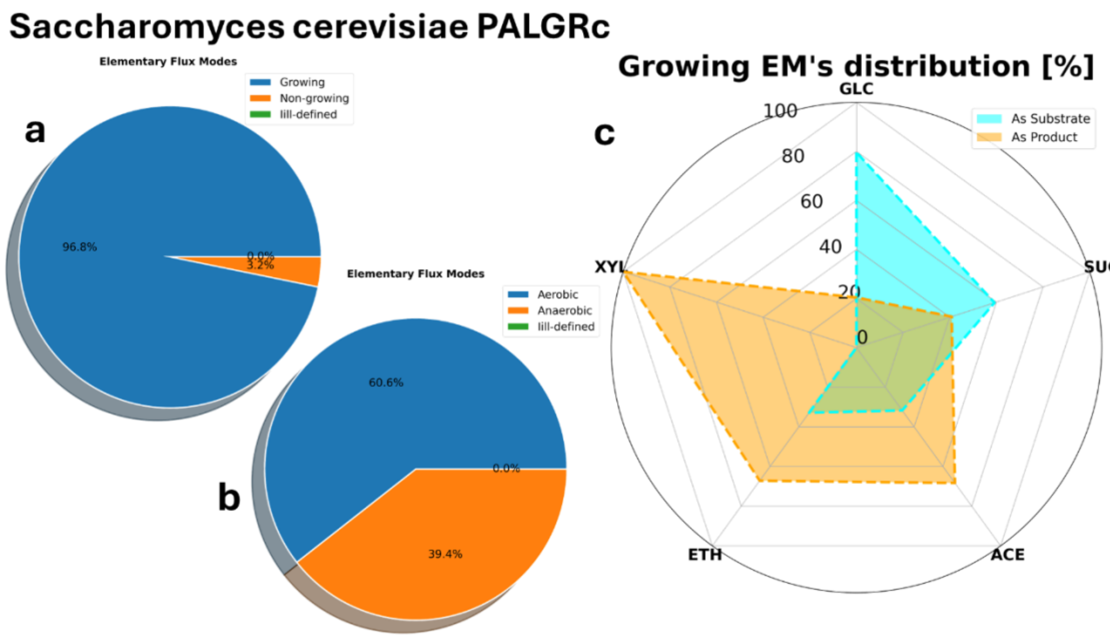


**Supplementary Figure 4** a) Distribution of EMs under the growth constraint for strain SAC*. b) Distribution of EMs under the constraint of Aerobiosis. c) Distribution of EMs under the constraints of single substrate consumption and principal produced compounds.

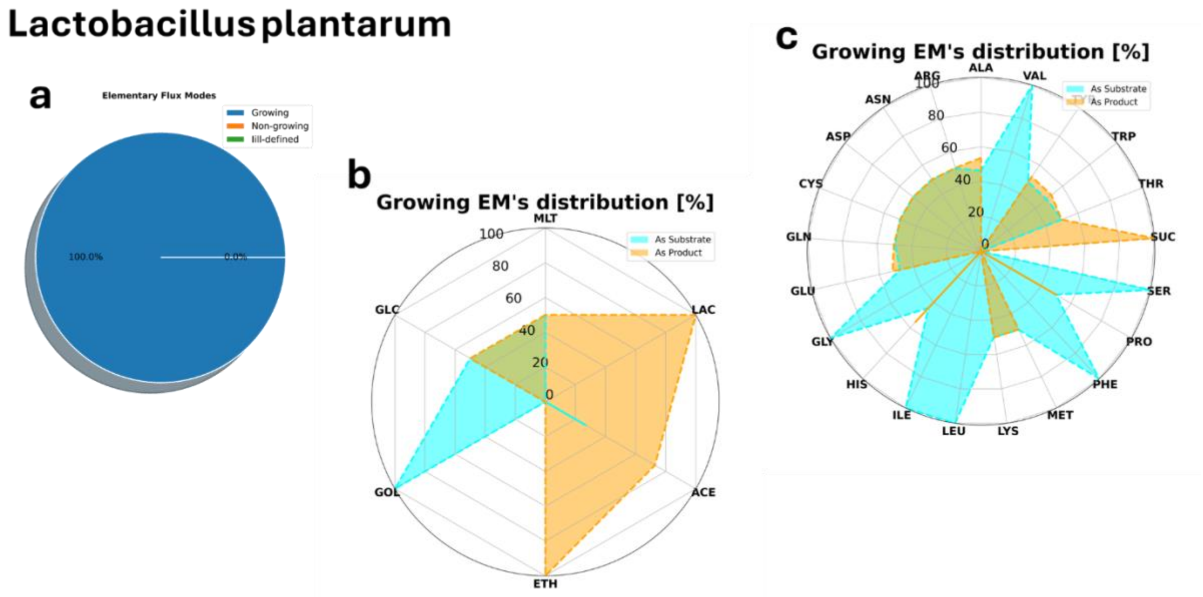


**Supplementary Figure 5** a) Distribution of EMs under the growth constraint for strain LAB. b) Distribution of EMs under the constraints of single substrate consumption and principal produced compounds. c) Distribution of EMs under the constraints of single substrate consumption and principal produced amino acids.

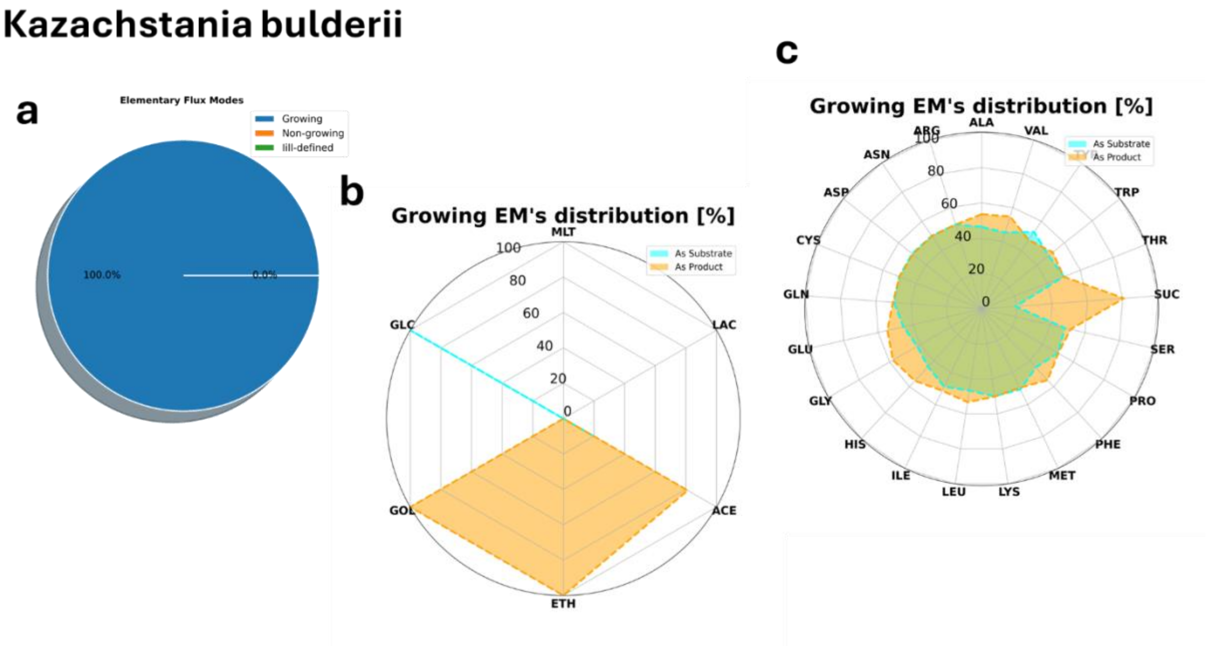


**Supplementary Figure 6** a) Distribution of EMs under the growth constraint for strain LAB. b) Distribution of EMs under the constraints of single substrate consumption and principal produced compounds. c) Distribution of EMs under the constraints of single substrate consumption and principal produced amino acids.

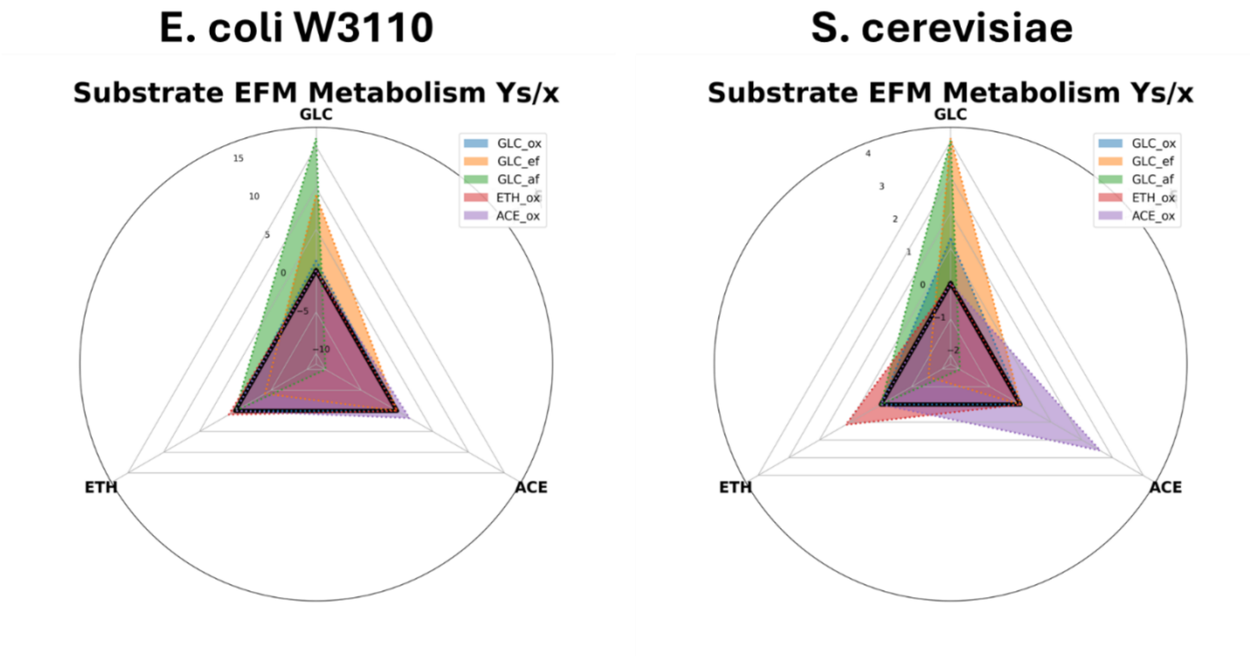


**Supplementary Figure 7** Selected active set for ECO (left) and SAC (right) corresponding to the oxidation of glucose, glucose fermentation to ethanol, glucose fermentation to acetic acid, ethanol oxidation and acetic acid oxidation for both organisms.

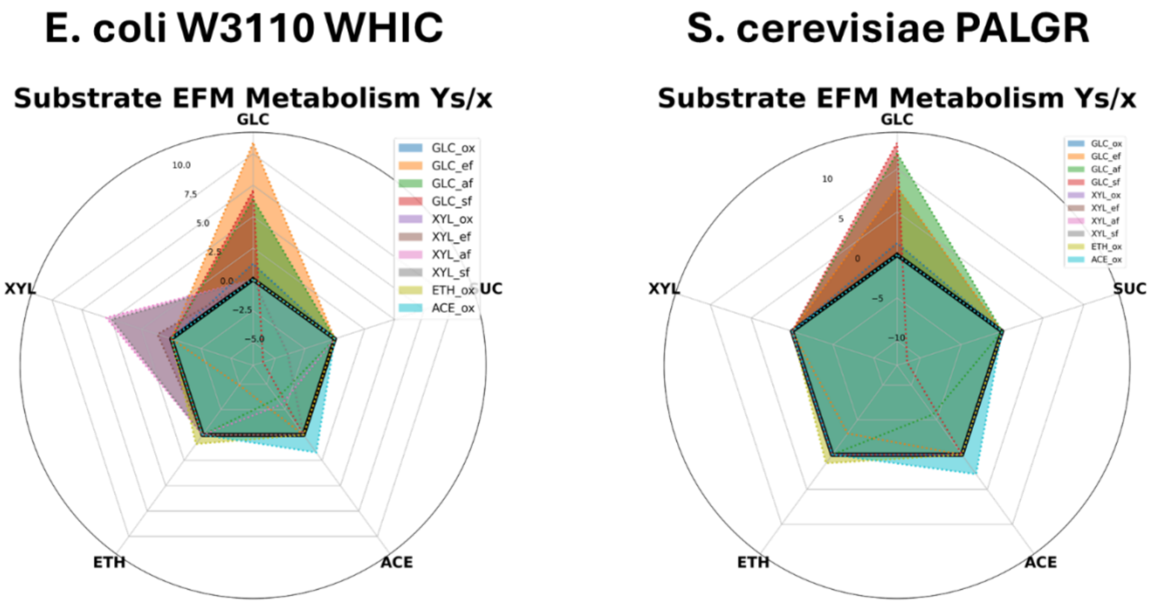


**Supplementary Figure 8** Selected active set for ECO* (left) and SAC* (right) corresponding to the oxidation of glucose, glucose fermentation to ethanol, glucose fermentation to acetic acid, oxidation of xylose, xylose fermentation to ethanol, xylose fermentation to acetic acid, ethanol oxidation and acetic acid oxidation for both organisms. SAC* xylose related EMs are found always as 0 valued vectors.

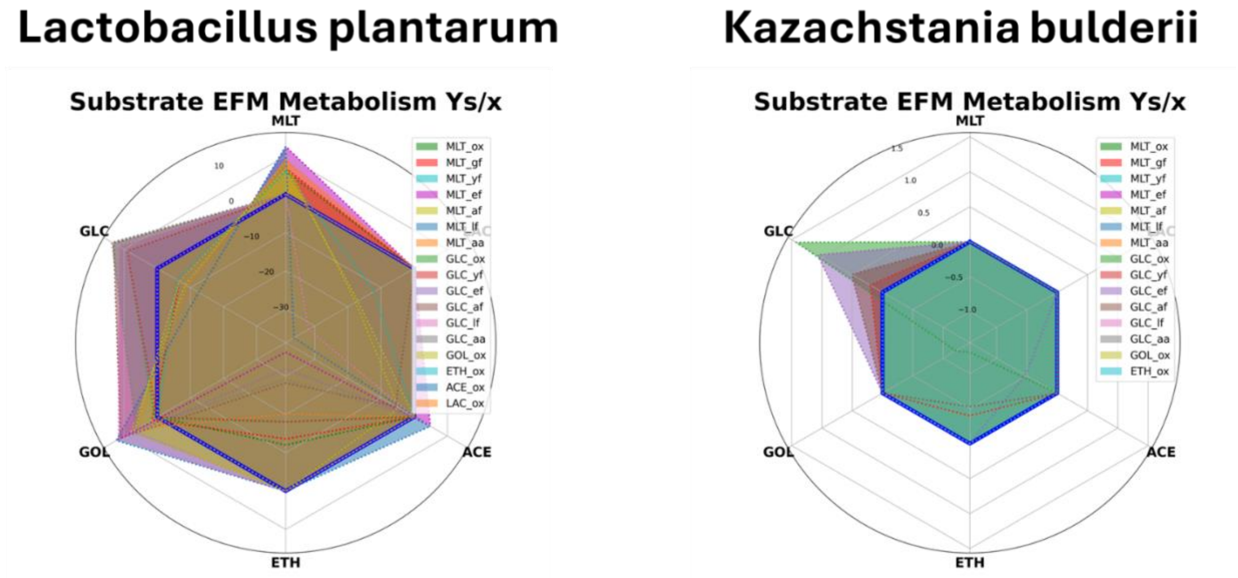


**Supplementary Figure 9** Sets of EMs as principal substrates for LAB (left) and KAZ (right). For these strains, no single substrate EMs were found, for LAB it is because the auxotrophies found for certain amino acids, while for Kazachstania bulderi, the interruption of the cytochromes results in a faulty oxidation reaction chain and therefore an energy imbalance that is solved by the co-utilization of metabolites for TCA replenishment such as succinate or others.

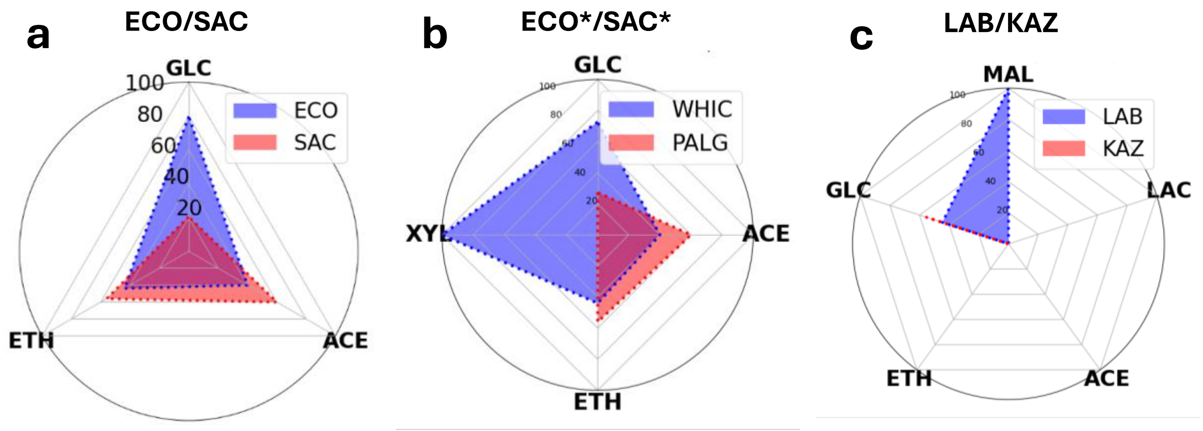


**Supplementary Figure 10** Metabolic Niche Advantage or aggressivity of each microorganism in comparison with each co-culture pair a) ECO/SAC pair, b) ECO*/SAC* pair, and c) LAB/KAZ pair

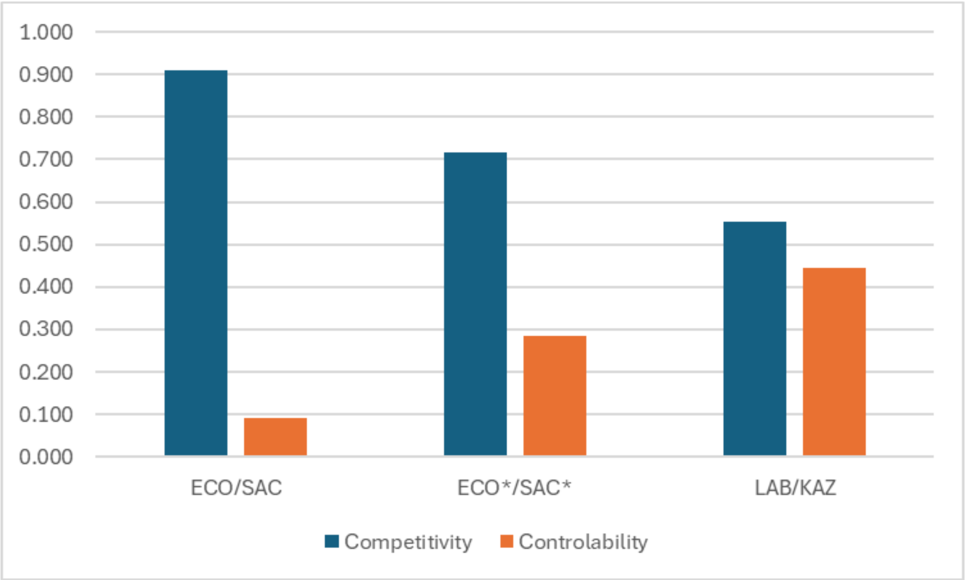


**Supplementary Figure 11**General Competitivity and Controllability of the three pairs of microorganisms according to the selected active sets, LAB/KAZ yields were corrected by adding the yield amount necessary of GLC to consume and overproduce the amino acids necessary for LAB growth.

**Supplementary Table 1** Extreme Yields from EMs, Aggressivity, Competitivity in all substrates as well as Global Competitivity and Controllability calculated for the pair ECO/SAC

| in mol/mol | Y GLC | Y ETH | Y ACE |
|---|---|---|---|
| ECO | -115.35 | -25.33 | -37.50 |
| SAC | -29.50 | -32.28 | -56.20 |
| Agressivity | Y GLC | Y ETH | Y ACE |
| ECO | 0.796 | 0.440 | 0.400 |
| SAC | 0.204 | 0.560 | 0.600 |
| Competitivity | GLC | ETH | ACE |
| θi | 0.407 | 0.879 | 0.800 |
| GLOBAL θ | Controlability | | |
| 0.909 | 0.091 | | |

**Supplementary Table 2** Extreme Yields from EMs, Aggressivity, Competitivity in all substrates as well as Global Competitivity and Controllability calculated for the pair ECO*/SAC*

| in mol/mol | Y GLC | Y XYL | Y ETH | Y ACE |
|---|---|---|---|---|
| ECO* | -78.42631267 | -162.465115 | -25.3332807 | -37.5019 |
| SAC* | -29.50166667 | 0 | -32.2846099 | -56.1991 |
| Agressivity | Y GLC | Y XYL | Y ETH | Y ACE |
| ECO* | 0.726654137 | 1 | 0.439677336 | 0.400229 |
| SAC* | 0.273345863 | 0 | 0.560322664 | 0.599771 |
| Competitivity | GLC | XYL | ETH | ACE |
| θi | 0.546691726 | 0 | 0.879354673 | 0.800458 |
| GLOBAL θ | Controlability | | | |
| 0.71671968 | 0.28328032 | | | |

**Supplementary Table 3** Extreme Yields from EMs, Aggressivity, Competitivity in all substrates as well as Global Competitivity and Controllability calculated for the pair LAB/KAZ. ** Corrected for the Substrate consumption needs for amino acid excretion

| in mol/mol | MLT | GLC | MLT** | GLC** |
|---|---|---|---|---|
| LAB | -12.494 | -13.64 | -12.494 | -13.64 |
| KAZ | 0 | -1.42 | 0 | -16.9369 |
| Agressivity | MLT | GLC | MLT** | GLC** |
| LAB | 1 | 0.905710491 | 1 | 0.446088 |
| KAZ | 0 | 0.094289509 | 0 | 0.553912 |
| Competitivity | MLT | GLC | MLT** | GLC** |
| θi | 0 | 0.188579017 | 0 | 0.892176 |
| GLOBAL θ | Controlability | | | |
| 0.553911797 | 0.446088203 | | | |

### Supplementary Note #3: MONCKS hybrid cybernetic modelling

In this work we used a cybernetic framework based on the works of Kompala [1], Ramkrishna [2], Varner [3], Song [4], Ramkrishna [5] and in our previous work adapting this framework to co-culture simulations called MONCKS and published in Martinez *et al.* [6]. In MONCKS each microorganism is modelled by a set of Monod type ODEs that are used as kernel sections for the interaction to a common environmental model 'The reactor' which will at any given time calculate its material balance to generate the variables the microorganisms will evaluate to set their internal regulation variables (cybernetic variables) and therefore compute their growth, consumption, and production. The summatory of the effects of all participant cells each time delta then is used in the reactor's next material balance. In this work we have implemented MONCKS to a more general computational toolbox published as BioMaSS: Bioprocess Modelling and Simulation Software, available in the Gitlab repository at https://gitlab.uliege.be/mipi/published-software/mbiomas-core. This latter toolbox is capable of reading Model descriptions and processes conditions in csv. files to generate the diverse microorganism behaviour vectors and reactor or environmental sections to then extend de simulation by Euler numerical approximations.

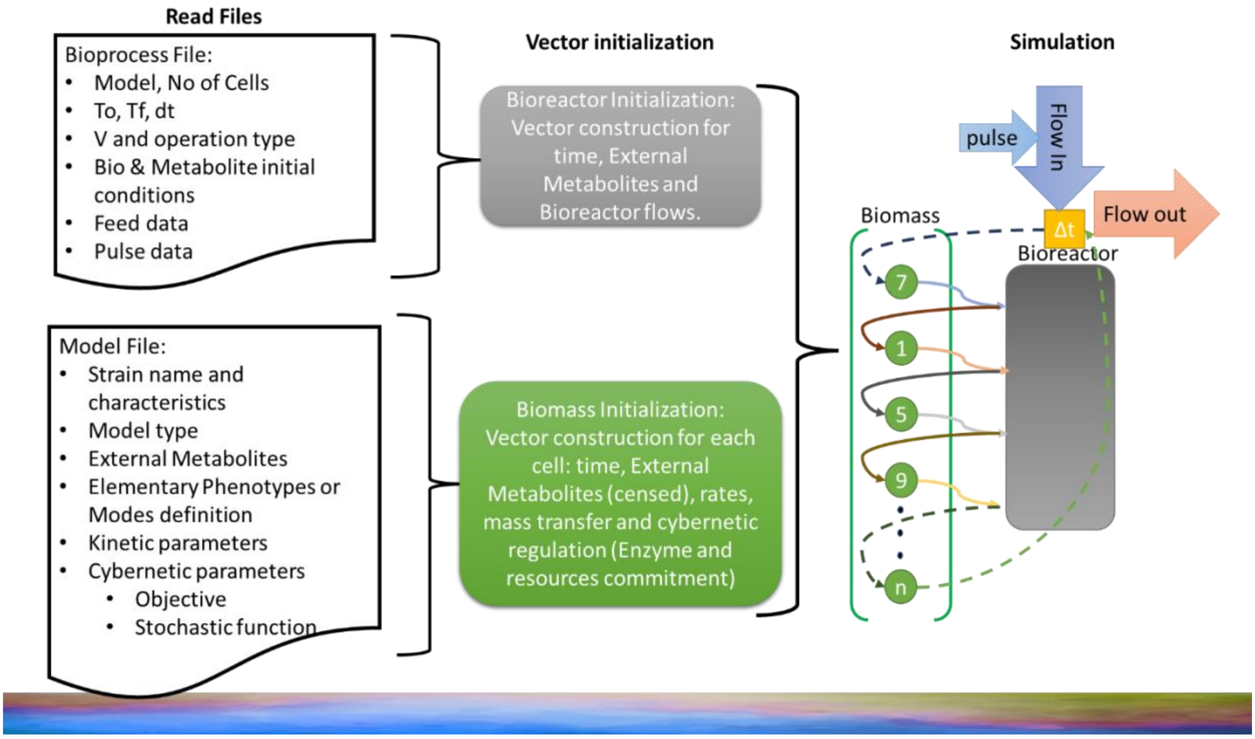


**Supplementary Figure 12** Main diagram representing the simulation strategy utilized in this work for MONCKS approach. Model files for each strain are read and transformed into vectors containing the internal fluxes, rates and regulation variables. Bioprocess or Environmental File is read to address the initial conditions that will determine the biomass behaviour and the external fluxes, pulses, dilution rates, derived from external environment perturbations. After this initial process the simulation calculates by Euler approximations the material balance in the environment and the effects of the biomass according to their growth, consumption, and production. In the case of this work the cellular effective rates were calculated based on a hybrid cybernetic model architecture with the EMs active set selected from the metabolic networks.

In this work the cellular effective rates were calculated based on a hybrid cybernetic model architecture with the EMs active set selected from the metabolic Networks described in the **Supplementary Note 2.** This active EMs yields then were used to calculate the consumption and production rates of each strain regulated by the cybernetic variables calculated in base of the metabolic objective of growth rate. A simplification of the information transfer can be found in the **Supplementary Figure 13**, and a detail information of the cybernetic mathematical approach can be found in Martinez *et al.* [6].

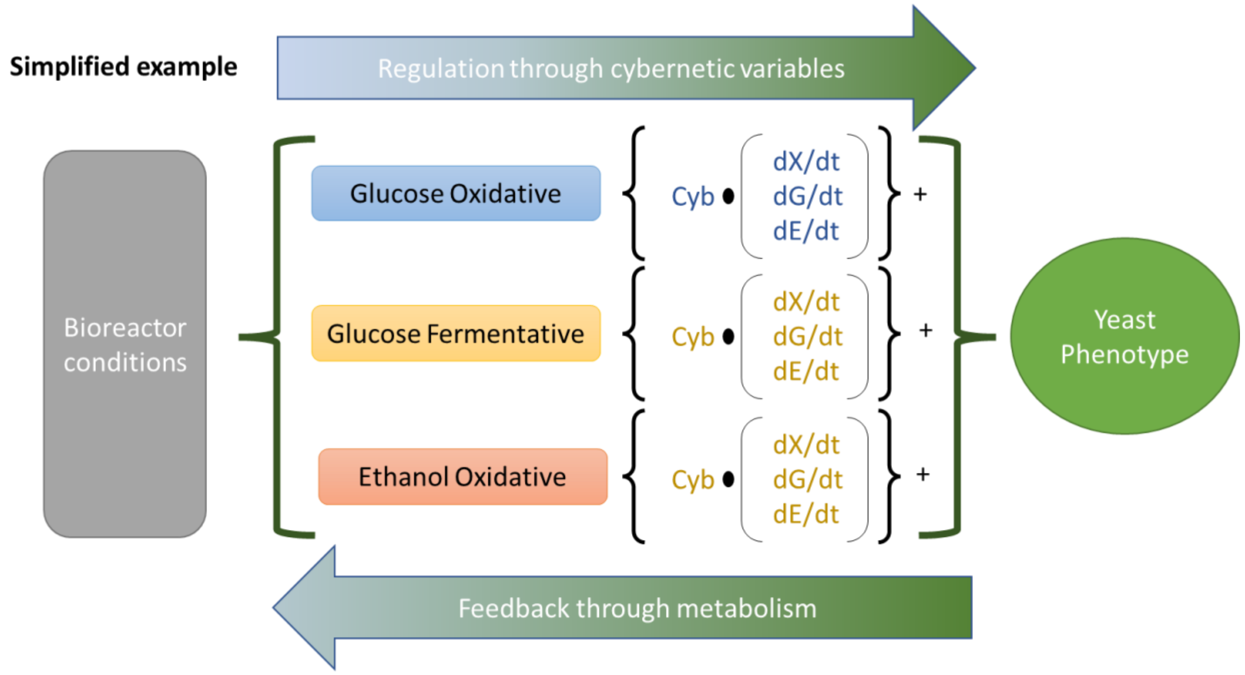


**Supplementary Figure 13** Information transfer between an example biomass, e.g. a yeast and the environmental or bioreactor conditions. The latter ones determine the capabilities of each Metabolic state represented by a single EMs in terms of a metabolic objective, e.g. growth. The latter set the cybernetic variables that regulate the Monod type growth, consumption, and production rates. With all the latter and the Biomass then the feedback on the environmental conditions is calculated in terms of instantaneous volumetric rates. In this way the system can be simulated every time delta with the help of Euler numerical approximations.

The EMs were used as found in the elementary mode analysis, except for the ECO fermentative production of acetate and ethanol from glucose, which was severely overestimated by the highest productive EMs, therefore a similar EMs was selected as fermentation but producing other byproducts were selected to be combined and fit the experimental yields found in ECO axenic cultures. This resulted in a similar EM yield composition but with a ten times reduction in the acetate production yield. The final EMs used for each strain, ECO, SAC, ECO* and SAC* were then transformed from Yields based on mol as in the stoichiometry reaction networks to g/g based on the formula weights of each metabolite and the summatory of the formula weights times the stoichiometry of the biomass reaction as an estimation for the biomass formula weight in the model. The final used yield values for the models can be found in the following tables (**Supplementary Tables 4-7**).

**Supplementary Table 4** Yields for the external metabolites used for the ECO models. Yields in g of metabolite/g of biomass

| Escherichia coli W3110 wt. (ECO) | Metabolite | Gox | Gef | Gaf | Eox | Aox |
|---|---|---|---|---|---|---|
| | **GLC** | -1.371 | -4.4 | -4.326 | 0 | 0 |
| | **ETH** | 0 | 1.663 | 0 | -1.231 | 0 |
| | **ACE** | 0 | 0 | 2.118 | 0 | -2.794 |

**Supplementary Table 5** Yields for the external metabolites used for the SAC models. Yields in g of metabolite/g of biomass

| Saccharomyces cerevisiae CENPK pchi-eGFP (SAC) | Metabolite | Gox | Gef | Gaf | Eox | Aox |
|---|---|---|---|---|---|---|
| | **GLC** | -1.236 | -9.112 | -16.01 | 0 | 0 |
| | **ETH** | 0 | 4.168 | 0 | -0.899 | 0 |
| | **ACE** | 0 | 0 | 10 | 0 | -1.734 |

**Supplementary Table 6** Yields for the external metabolites used for the ECO* models. Yields in g of metabolite/g of biomass. For simulations the plasmids were not considered since then the active selection set needed to be modified based on yield experimental data, but in this work the predictive nature of the central metabolism effects on growth and environment was preferred.

| Escherichia coli WHIC | Metabolite | Gox | Gef | Gaf | Xox | Gef | Xaf | Eox | Aox |
|---|---|---|---|---|---|---|---|---|---|
| | **GLC** | -1.236 | -10.879 | -6.4 | 0 | 0 | 0 | 0 | 0 |
| | **XYL** | 0 | 0 | 0 | -1.161 | -18.781 | -5.463 | 0 | 0 |
| | **TYR** | 0 | 0 | 0 | 0 | 0 | 0 | 0 | 0 |
| | **CMA** | 0 | 0 | 0 | 0 | 0 | 0 | 0 | 0 |
| | **GOL** | 0 | 0 | 0 | 0 | 0 | 0 | 0 | 0 |
| | **ETH** | 0 | 4.929 | 0 | 0 | 9.113 | 0 | -0.899 | 0 |
| | **ACE** | 0 | 0 | 3.599 | 0 | 0 | 3 | 0 | -1.734 |
| | **SUC** | 0 | 0 | 0 | 0 | 0 | 0 | 0 | 0 |
| | **CO2** | 0.429 | 4.36 | 0.002 | 0.319 | 3.143 | 0.028 | 0.334 | 1.159 |

**Supplementary Table 7** Yields for the external metabolites used for the SAC* models. Yields in g of metabolite/g of biomass. For simulations the plasmids were not considered since then the active selection set needed to be modified based on yield experimental data, but in this work the predictive nature of the central metabolism effects on growth and environment was preferred.

| | Metabolite | Gox | Gef | Gaf | Eox | Aox |
|---|---|---|---|---|---|---|
| Saccharomyces cerevisiae PTA 408 (SAC*) | **GLC** | -1.371 | -4.4 | -4.326 | 0 | 0 |
| | **XYL** | 0 | 0 | 0 | 0 | 0 |
| | **TYR** | 0 | 0 | 0 | 0 | 0 |
| | **CMA** | 0 | 0 | 0 | 0 | 0 |
| | **GOL** | 0 | 0 | 0 | 0 | 0 |
| | **ETH** | 0 | 1.663 | 0 | -1.231 | 0 |
| | **ACE** | 0 | 0 | 2.118 | 0 | -2.794 |
| | **SUC** | 0 | 0 | 0 | 0 | 0 |
| | **CO2** | 0.405 | 1.669 | 0.328 | 0.748 | 2.491 |

With this EMs Yields we estimated the maximum growth rates and the Monod saturation constants for each Metabolic state, e.g. glucose oxidation (Gox), glucose fermentation to ethanol (Gef), glucose fermentation to acetic acid (Gaf), ethanol oxidation (Eox) and acetate oxidation (Aox) for ECO via approximation to the behaviour of axenic batch cultures. The latter was performed for ECO and SAC strains. Approximation was performed by a Montecarlo and Metropolis hasting algorithms. After this parameter approximation, chemostat and AAMN cultures were used as validation for the models. After validation these parameters were used as estimations for the rates of the ECO* and SAC* simulations with different perturbations made to understand the capabilities of stabilization and control. In this case first XYL and GLC metabolic niches were given similar fitness performance by setting their rates the same, then a reduction of the GLC derived metabolism was tested by setting the rates growth on GLC reduced by a factor of 10. Chemostat simulations resulted in a slight increase in the SAC* population at equilibrium but still with ECO* domination. While in AAMN a small stabilization time is obtained but then control is lost by the increase of other uncontrolled metabolic niches such as acetate and ethanol. For that reason, we decided to limit the apparition of alternative uncontrolled niches by reducing the by-product production by q factor of 2 and a factor of 10. Similarly, chemostat co-cultures resulted in ECO* domination. Conversely, AAMN had increased control capabilities and in the case of by product yield reduction by a factor of 10 a clear 50/50 population ration between ECO*/SAC* was obtained. All files and pipelines used for the calculation of all simulations, axenic approximations and co-culture validations and predictive simulations can be found at https://gitlab.uliege.be/mipi/published-software/2024-cocultures the Gitlab repository. Below in this Supplementary Note we will present figures representing the simulations performed.

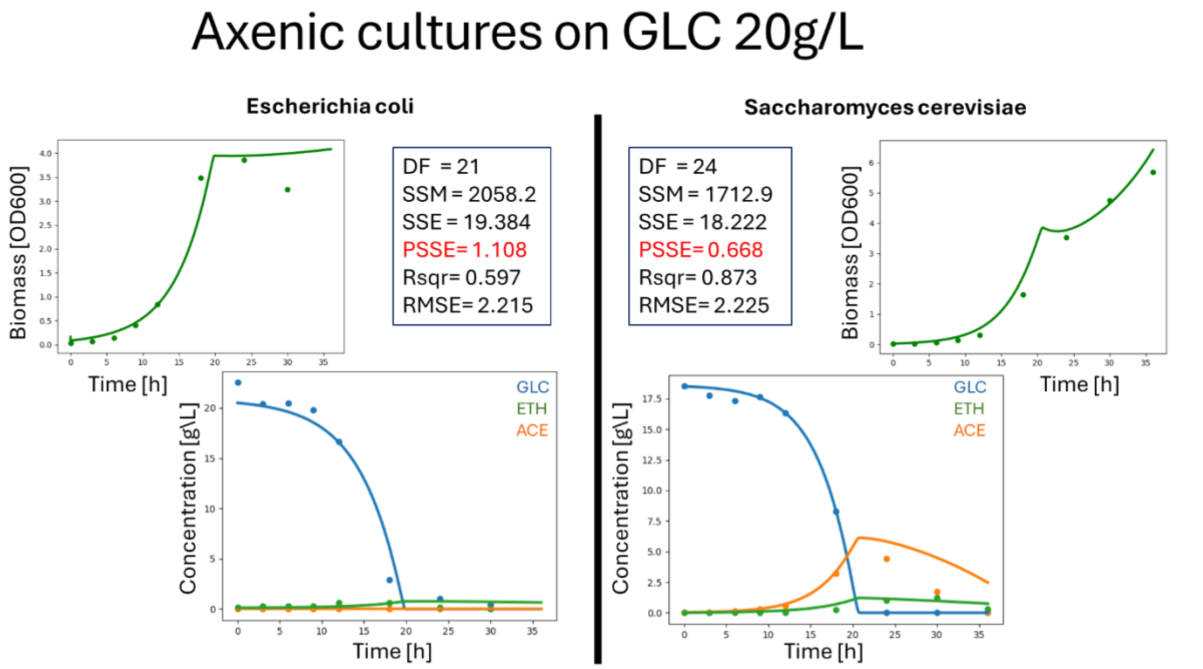


**Supplementary Figure 14** Experimental profiles (dots) for Biomass growth and principal metabolites for the axenic cultures of ECO (left) and SAC (right), and the simulated results (lines), as well as the values for parameter approximations. DF: degrees of freedom; SSM: Sum of Square Measurements; SSE: Sum of square Error; PSSE: Percentual Sum of Square Error; Rsqr: Pearson Correlation Coefficient; RMSE: Root Mean square Error.

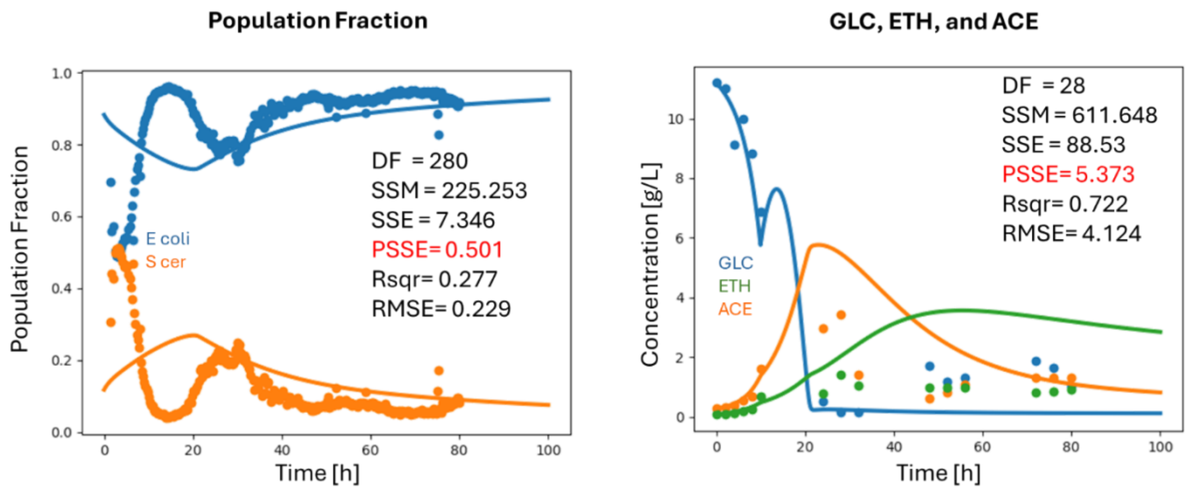


**Supplementary Figure 15** Experimental and model behaviour comparison. Population fraction (right) and External Metabolite composition (Left) as validation of the parameters found with the axenic cultures. As well as the values for parameter approximations. DF: degrees of freedom; SSM: Sum of Square Measurements; SSE: Sum of square Error; PSSE: Percentual Sum of Square Error; Rsqr: Pearson Correlation Coefficient; RMSE: Root Mean square Error.

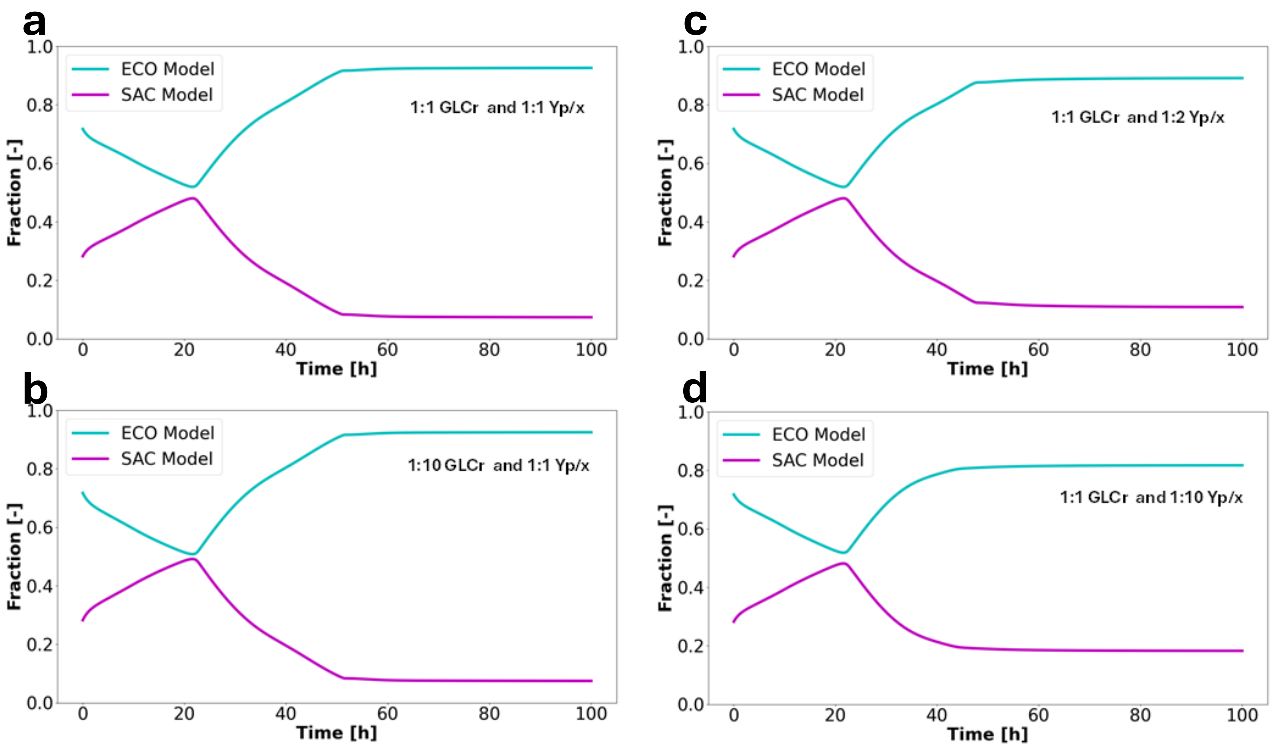


**Supplementary Figure 16** Chemostat Population fraction profiles for the simulation of the ECO*/SAC* mixtures a) simulation with GLC rates and XLY rates with equal values, b) simulation with GLC rates reduced by ten times in comparison to XYL, c) reduction of the byproduct production by half, d) reduction of the byproduct production by ten times.

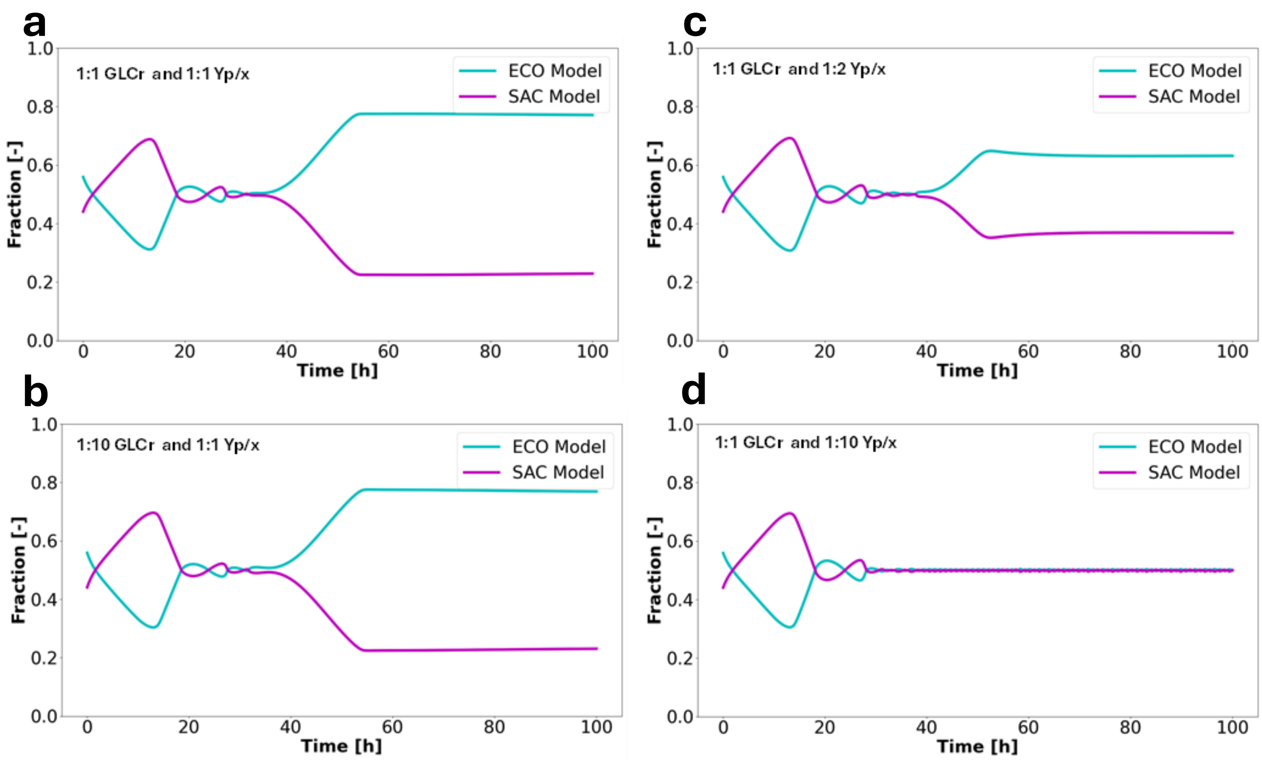


**Supplementary Figure 17** AAMN Population fraction profiles for the simulation of the ECO*/SAC* mixtures a) simulation with GLC rates and XLY rates with equal values, b) simulation with GLC rates reduced by ten times in comparison to XYL, c) reduction of the byproduct production by half, d) reduction of the byproduct production by ten times.

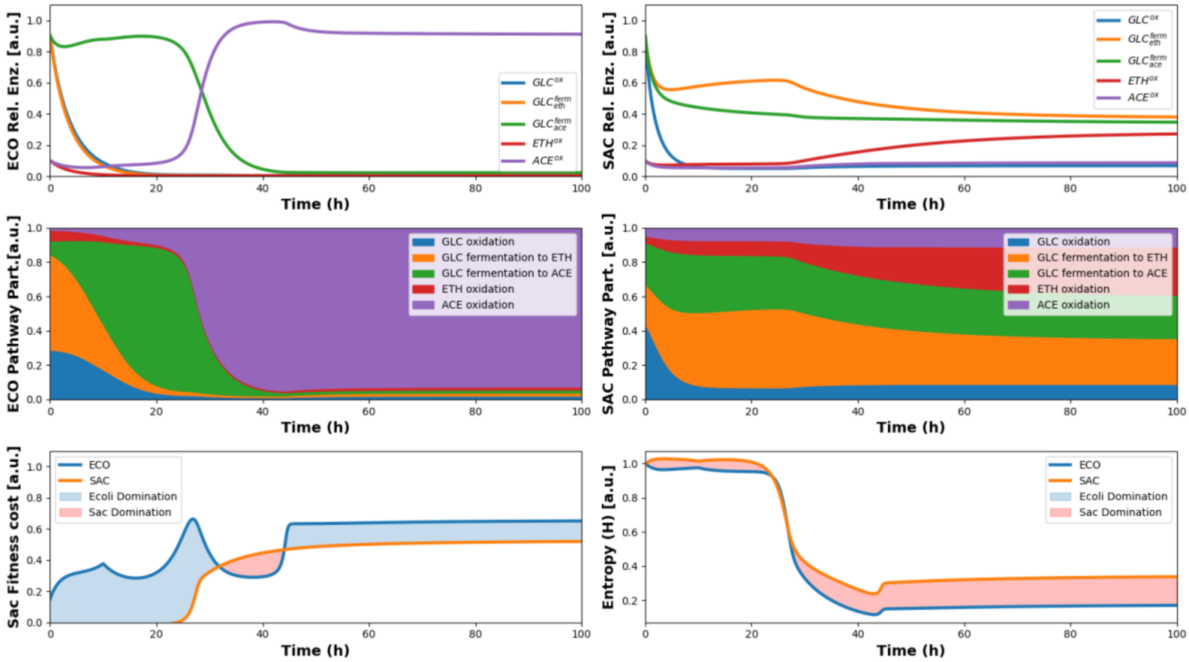


**Supplementary Figure 18** Simulated Cybernetic regulation of metabolism for the ECO/SAC in a chemostat co-culture.

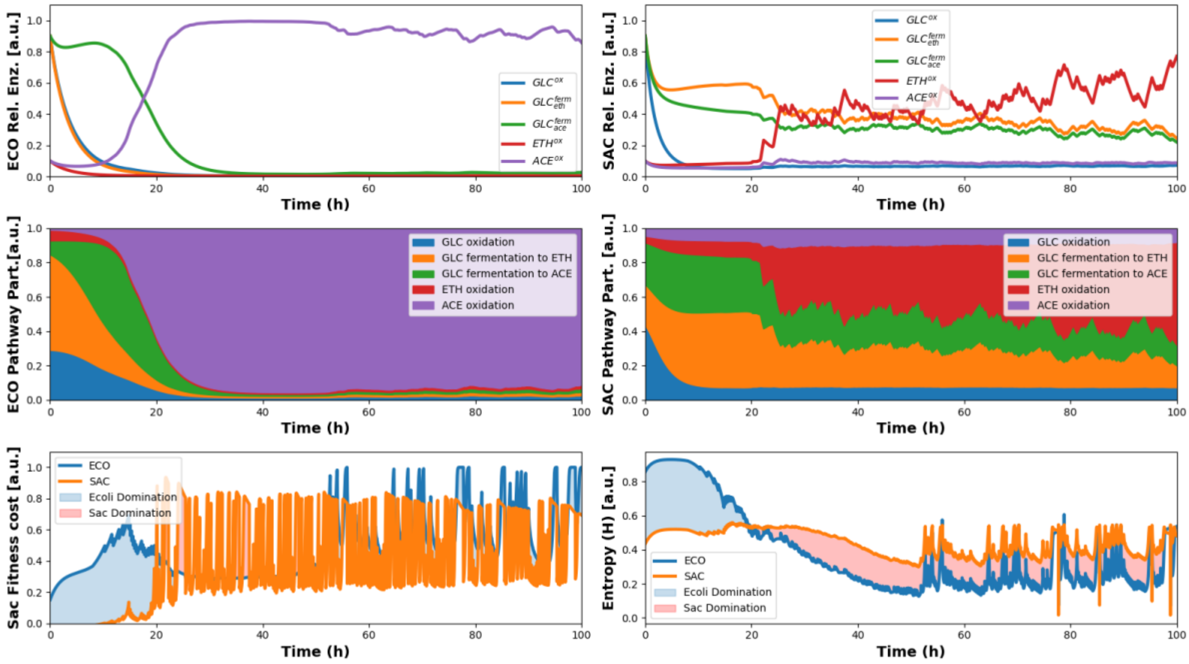


**Supplementary Figure 19** Simulated cybernetic regulation of metabolism for the ECO/SAC in a random pulse continuous co-culture.

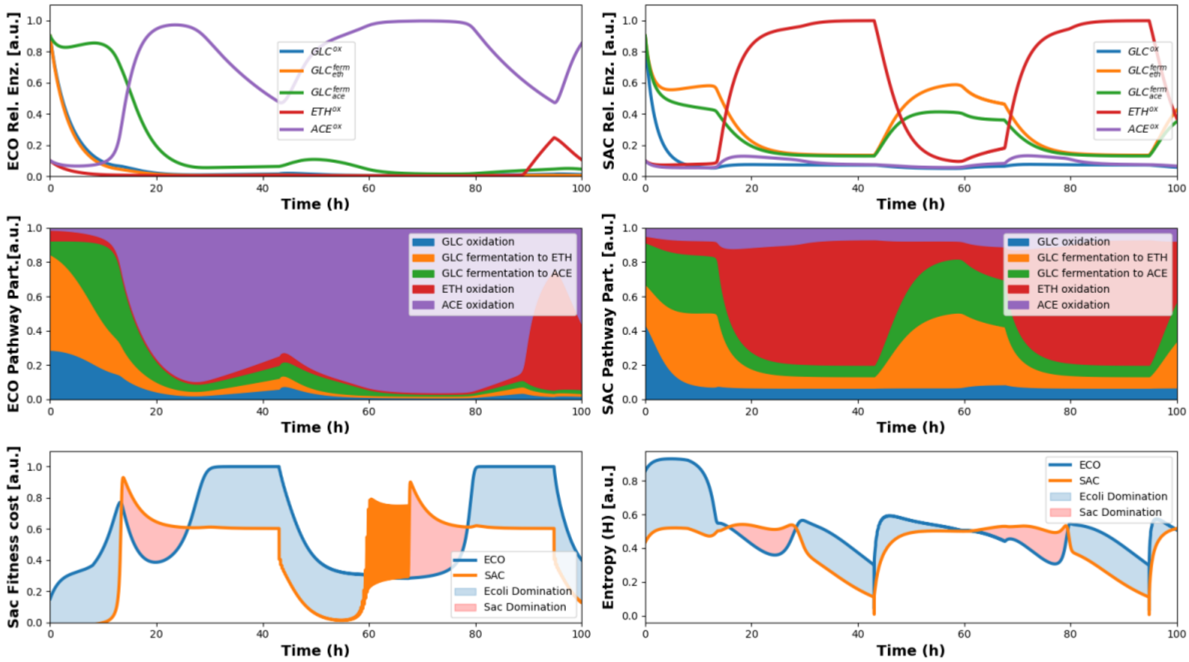


**Supplementary Figure 20** Simulated cybernetic regulation of metabolism for the ECO/SAC in a AAMN controlled co-culture.

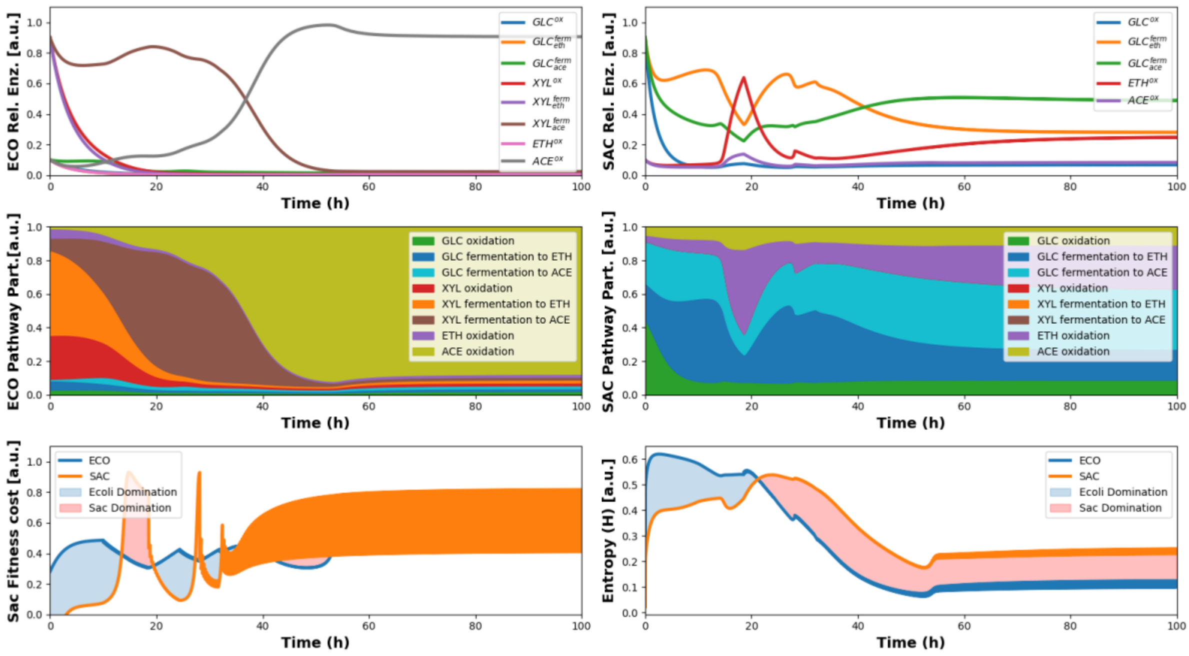


**Supplementary Figure 21** Simulated cybernetic regulation of metabolism for the ECO*/SAC* in a AAMN co-culture, parameters 1:1 for rates and yields.

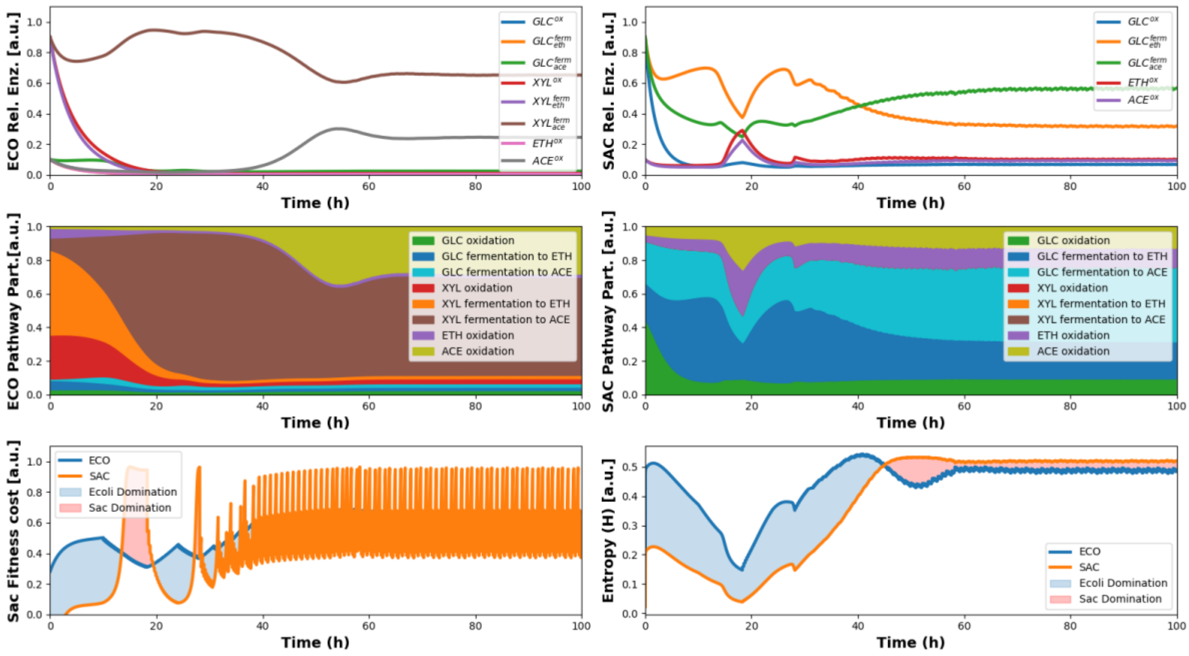


**Supplementary Figure 22** Simulated cybernetic regulation of metabolism for the ECO*/SAC* in a AAMN co-culture, parameters 1:1 for rates and 1:10 reduction in ACE and ETH yields.

#### *Supplementary Note 3 references:*

#### Supplementary Note #4: Shannon entropy as an index for competitivity and controllability.

In this work we define competitivity as the capability of the strains to allocate certain substrate into their metabolic networks to grow. This can be calculated from the individual Volumetric consumption rates from each individual organisms and comparing it to the total consumption rate. To make this comparison we decided to use the Shannon's entropy from information theory, which allows us to obtain an index in bits that tells us how much information is needed to describe the total number of states of a system and their composition. In other words, it is useful to attest the distribution of a system, and therefore is a good indicator for the task of estimating the probability of the distribution pulses or environmental changes will have in the environment. The mathematical formulas are described in material and methods, however here we want to present tables and graphs that support the behaviour of the indexes. First, we suppose a single substrate and a probability of being consumed by an organism A (Pa), the Probability of being consumed by B (Pb) would be therefore 1 – Pa so that Pa + Pb = 1 then we can calculate the Shannon partial entropies (ha) and (hb) for each microorganism and then the controllability and competitivity parameters such as in **Supplementary Table 8** and observed in **Supplementary Figure 18**

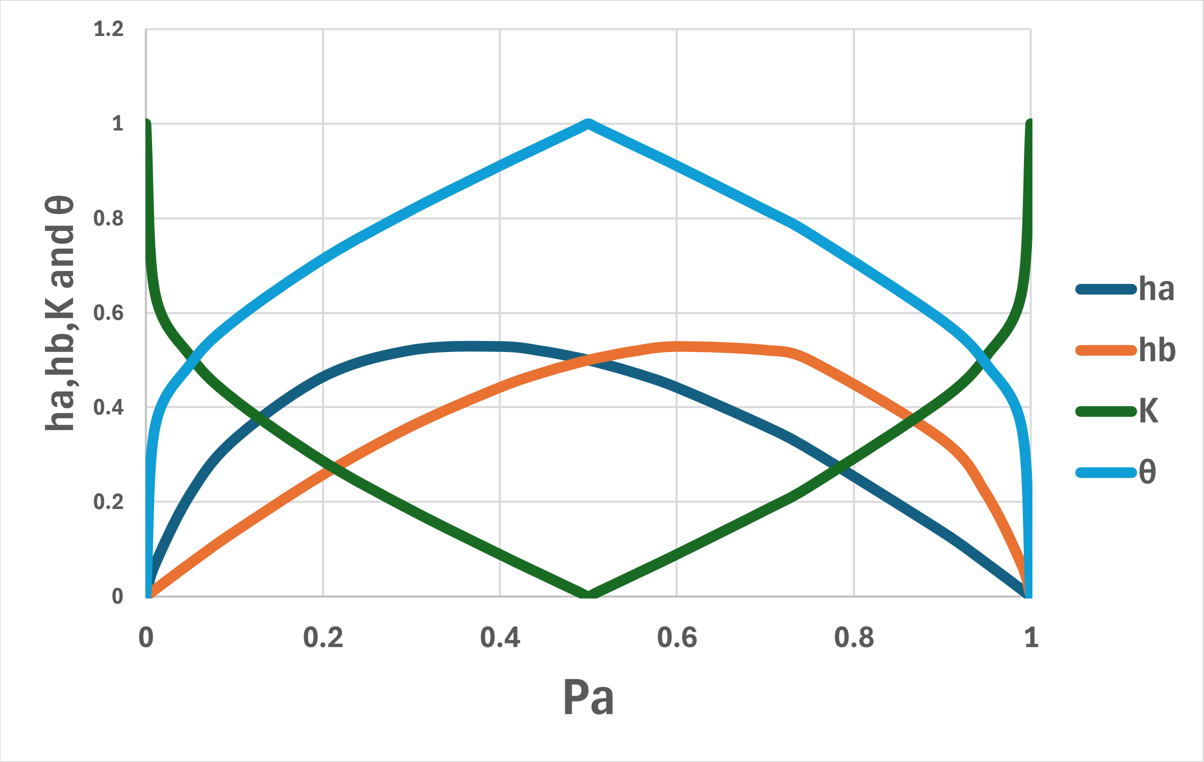


**Supplementary Figure 23** Behaviour of the partial entropies ha and hb corresponding to the microorganism A and B, as well as the indexes of controllability K and competitivity θ for the consumption of the substrate s in terms of the probability of it being consumed by A

**Supplementary Table 8** Table containing example values for the calculation of the Controllability and Competitivity of the system in a single substrate between two microorganisms A and B which at a given time their behaviour on the environment records the volumetric values of Qs.

| Probability of A (Qs of A/Qs global) Pa | Probability of B (Qs of B/Qs global) Pb | P.Entropy A -Pa log2 (Pa) ha | P. Entropy B -Pb log2 (Pb) hb | Controlability K | Competitivity θ |
|---|---|---|---|---|---|
| 1 | 0 | 0 | 0 | 1 | 0 |
| 0.990 | 0.010 | 0.014 | 0.066 | 0.645 | 0.355 |
| 0.950 | 0.050 | 0.070 | 0.216 | 0.509 | 0.491 |
| 0.900 | 0.100 | 0.137 | 0.332 | 0.417 | 0.583 |
| 0.750 | 0.250 | 0.311 | 0.500 | 0.233 | 0.767 |
| 0.700 | 0.300 | 0.360 | 0.521 | 0.183 | 0.817 |
| 0.600 | 0.400 | 0.442 | 0.529 | 0.089 | 0.911 |
| 0.550 | 0.450 | 0.474 | 0.518 | 0.044 | 0.956 |
| 0.510 | 0.490 | 0.495 | 0.504 | 0.009 | 0.991 |
| 0.500 | 0.500 | 0.500 | 0.500 | 0.000 | 1.000 |
| 0.490 | 0.510 | 0.504 | 0.495 | 0.009 | 0.991 |
| 0.450 | 0.550 | 0.518 | 0.474 | 0.044 | 0.956 |
| 0.400 | 0.600 | 0.529 | 0.442 | 0.089 | 0.911 |
| 0.300 | 0.700 | 0.521 | 0.360 | 0.183 | 0.817 |
| 0.200 | 0.800 | 0.464 | 0.258 | 0.287 | 0.713 |
| 0.100 | 0.900 | 0.332 | 0.137 | 0.417 | 0.583 |
| 0.050 | 0.950 | 0.216 | 0.070 | 0.509 | 0.491 |
| 0.010 | 0.990 | 0.066 | 0.014 | 0.645 | 0.355 |
| 0.000 | 1.000 | 0.000 | 0.000 | 1.000 | 0.000 |

The controllability index tends to be 0 when the probability of this substrate s to be consumed by each microorganism pull is 0.5 and equal, therefore pulses of this substrate would have little to no effect on the population state of the system. Conversely the control ability tends to be 1 when an organism utilizes exclusively this substrate for growth, therefore changes in this substrate will have great effects on the population that consumes it. Effectively, in this extreme case then the non-consumer has no possibilities to grow and therefore it would be extinguished from the systems. Therefore, for achieving maximum controllability to sustain two populations the optimum requires at least two substrates with maximum controllability and with inverse probabilities on each substrate. However, this is not often possible, therefore equilibrium and system control are only effective by temporal transitions between substrate conditions that would give rise to temporal fitness accordingly to the probabilities of this temporal environments to be more beneficial to each co-culture member. To calculate the total controllability of the system we utilize the mean K calculated for each substrate present in the media. It is important to note that this index tells us a comparison of the probabilities of substrate consumption therefore but not the direction the system is going not the speed at which it is reaching equilibria, or the size of the pulse or modification needed to make a visible effect on the system. For that other analysis of consumption rates can be performed, or appropriate control rules can be defined for an AAMN bioprocess.

### Supplementary Note #5: Calculation of auto-aggregation

The calculation of auto-aggregation was performed in a similar manner as in the article of Verastegui *et al*. [1]. First, raw data was cleaned from noise by eliminating event with negative and cero values of all measured channels. Then doublets were eliminated by the ratio of FSC-A and FSC-H where selected above 2 were eliminated. Samples that after cleaning resulted in less than 20 000 events or with more than 10% of noise or doublets were re-examined and/or eliminated. Finally, auto-aggregation was performed by obtaining the mean FSC-A and standard deviation that describes a single cell population. In this case we calculated the Grand Mean and Standard deviations from all samples and assuming that mostly all of them are in a single cell state use this to select events in all samples with values of FSC-A greater than 3 times this standard deviation, therefore with a probability near 97% we can assume this are cells with different morphology than a single cell. This indicates that they can be either supersized or filamented cells or auto-aggerated cells. The increased values of SSC-A suggest that the latter is the most appropriate possibility since a hyper-rough surface given by several cells agglomerated could be responsible for this SSC-A, otherwise cells would need to increase intracellular complexity and that is less probable for the bacterial kinds. This is rather relevant for the present work control algorithm since it represents an error in the approximation of mass consuming the different metabolites pulsed for control, more about we cannot determine by FC the composition if this agglomerates, and therefore they represent a serious challenge for the controller and actuator as presented in this work. Further developments of control rules and FC analysis should be performed for allowing better stability and robustness to AAMN as presented in this work. Below we present the results for the calculation of the auto-aggregations found for the different experiments (**Supplementary Figure 24-27**).

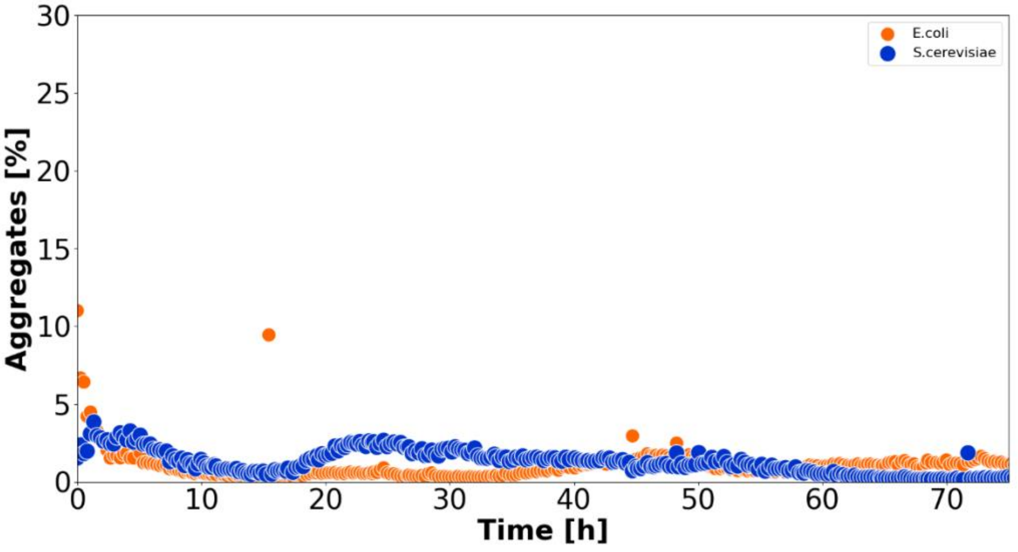


**Supplementary Figure 24** Auto-aggregation profiles for ECO and SAC for the chemostat experiments.

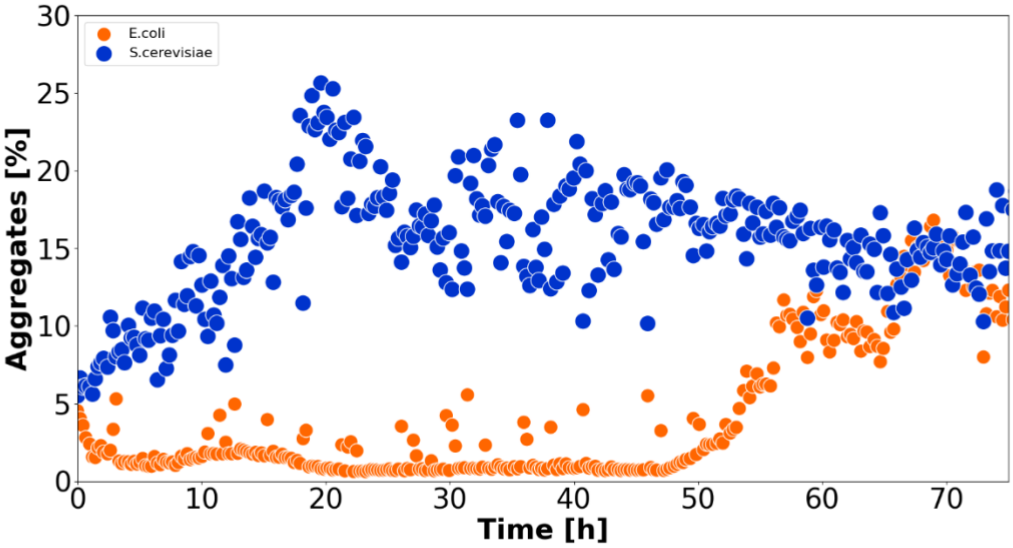


**Supplementary Figure 25** Auto-aggregate profiles for ECO and SAC for the AAMN experiments

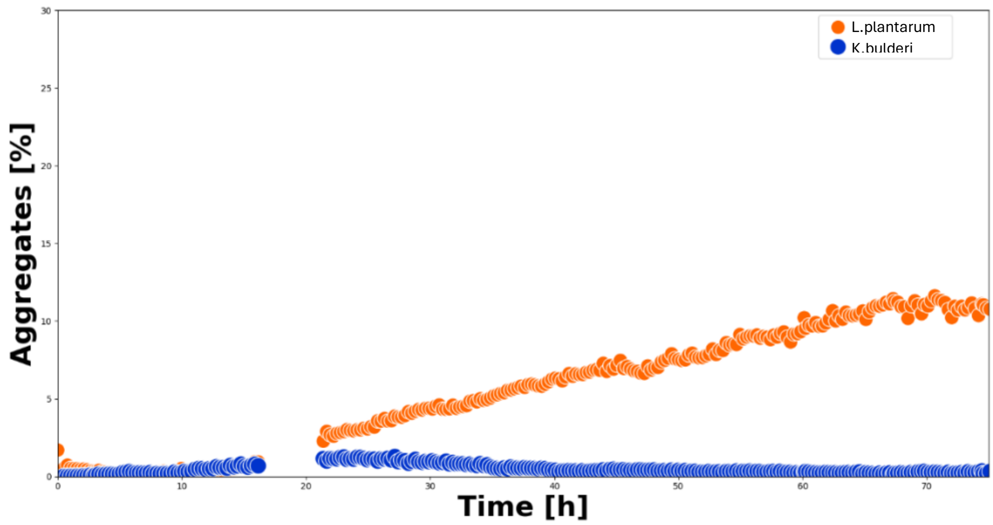


**Supplementary Figure 26** Auto-aggregate profiles for LAB and KAZ for the chemostat experiments

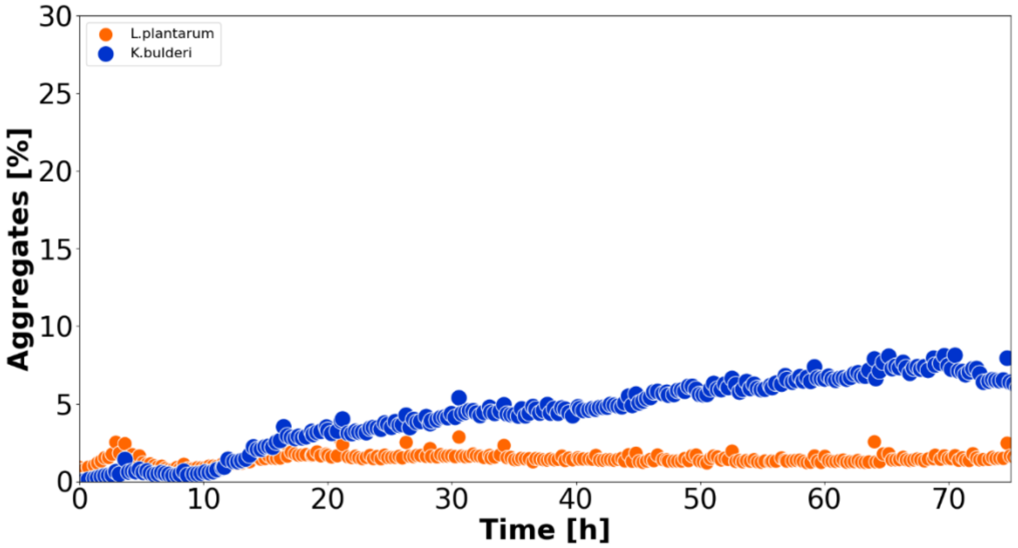


**Supplementary Figure 27** Auto-aggregate profiles for LAB and KAZ for the AAMN experiments

##### *Supplementary Note 5 references:*


1. Velastegui, E., Quezada, J., Guerrero, K. et al. Is heterogeneity in large-scale bioreactors a real problem in recombinant protein synthesis by Pichia pastoris?. Appl Microbiol Biotechnol 107, 2223–2233 (2023). https://doi.org/10.1007/s00253-023-12434-2

#### Supplementary Note #6: Media preparation and analysis specifications

In this work we decided to work with minimal mineral media, to avoid undesired or unquantifiable interactions and effects through experiments and modelling. However, this was not always possible due to some auxotrophies and feed requirements for some organisms. That is mainly the case the LAB strain which has several amino acid auxotrophies. The preparation of media is described in the **Methods** section, however, here we present the tables of media composition. Finally, we also attach the specific HPLC mobile phase media composition used for the analysis of media composition across all experiments.

**Supplementary Table 9** Verduyn media and adapted MRS formulations used for the cultivation of the different microbial strains was developed according to requirements found in the literature.

**Verduyn media**

| Components | Concentration | Units |
|---|---|---|
| $KH_2PO_4*H_2O$ | 9,34 | g/L |
| $K_2HPO_4*7H_2O$ | 6,309 | g/L |
| Ammonium sulfate | 5 | g/L |
| Magnesium sulfateheptahydrate | 0,5 | g/L |
| Ethylenediaminetetraacetic acid (EDTA) | 95,55 | mg/L |
| Zinc sulfate heptahydrate | 22,5 | mg/L |
| Manganese(II) chloride tetrahydrate | 5 | mg/L |
| Cobalt(II) chloride heptahydrate | 1,5 | mg/L |
| Copper(II) sulfate penthahydrate | 1,5 | mg/L |
| Sodium molybdate dihydrate | 2 | mg/L |
| Calcium chloride dihydrate | 22,5 | mg/L |
| Iron(II) sulfate heptahydrate | 15 | mg/L |
| Boric acid | 5 | mg/L |
| Potassium iodide | 0,5 | mg/L |
| D-biotin | 0,05 | mg/L |
| Calcium pantothenate | 1 | mg/L |
| Nicotinic acid | 1 | mg/L |
| myo-inositol | 25 | mg/L |
| Thiamine HCl | 1 | mg/L |
| Pyridoxine HCl (vit. B6) | 1 | mg/L |
| Para-aminobenzic acid | 0,2 | mg/L |
| Cobalamine | 0,5 | µg/L |
| Folic acid | 0,5 | µg/L |

**Adapted MRS Media**

| Components | Concentration | Units |
|---|---|---|
| Triptone | 10 | g/L |
| Yeast extract | 5 | g/L |
| Meat extract | 5 | g/L |
| $KH_2PO_4*H_2O$ | 12,68 | g/L |
| $K_2HPO_4*7H_2O$ | 2,18 | g/L |
| Ammonium sulfate | 5 | g/L |
| Monopotassium phosphate | 3 | g/L |
| Magnesium sulfateheptahydrate | 0,5 | g/L |
| Ethylenediaminetetraacetic acid (EDTA) | 95,55 | mg/L |
| Zinc sulfate heptahydrate | 22,5 | mg/L |
| Manganese(II) chloride tetrahydrate | 5 | mg/L |
| Cobalt(II) chloride heptahydrate | 1,5 | mg/L |
| Copper(II) sulfate penthahydrate | 1,5 | mg/L |
| Sodium molybdate dihydrate | 2 | mg/L |
| Calcium chloride dihydrate | 22,5 | mg/L |
| Iron(II) sulfate heptahydrate | 15 | mg/L |
| Boric acid | 5 | mg/L |
| Potassium iodide | 0,5 | mg/L |
| D-biotin | 0,05 | mg/L |
| Calcium pantothenate | 1 | mg/L |
| Nicotinic acid | 1 | mg/L |
| myo-inositol | 25 | mg/L |
| Thiamine HCl | 1 | mg/L |
| Pyridoxine HCl (vit. B6) | 1 | mg/L |
| Para-aminobenzic acid | 0,2 | mg/L |
| Cobalamine | 0,5 | µg/L |
| Folic acid | 0,5 | µg/L |

**Supplementary Table 10 :** UPLC conditions for the mobile phase.

| Time (minutes) | TFA in water (%) | TFA in methanol (%) |
|---:|---:|---:|
| 1 | 100 | 0 |
| 2 | 95 | 5 |
| 5.5 | 85 | 15 |
| 5.6 | 0 | 100 |
| 6.6 | 0 | 100 |
| 6.7 | 100 | 0 |
